# Evaluating noise correction approaches for non-invasive electrophysiology of the human spinal cord

**DOI:** 10.1101/2024.09.05.611423

**Authors:** Emma Bailey, Birgit Nierula, Tilman Stephani, Burkhard Maess, Vadim Nikulin, Falk Eippert

**Affiliations:** Max Planck Research Group Pain Perception, Max Planck Institute for Human Cognitive and Brain Sciences, Leipzig, Germany; Research Group Neural Interactions and Dynamics, Department of Neurology, Max Planck Institute for Human Cognitive and Brain Sciences, Leipzig, Germany; Methods and Development Group Brain Networks, Max Planck Institute for Human Cognitive and Brain Sciences, Leipzig, Germany

## Abstract

The spinal cord is a vital component of the central nervous system for the processing of sensorimotor information transmitted between the body and the brain. Electrospinography (ESG) is the most accessible non-invasive technique for recording spinal signals in humans, but the vast and detrimental impact of physiological noise (mostly of cardiac nature) has prevented widespread adoption. Here, we aim to address this issue by examining various denoising algorithms for cardiac artefact reduction – including principal component analysis-based techniques (PCA), independent component analysis-based approaches (ICA) and signal space projection (SSP). We observed that in situations where large numbers of spinal electrodes are used, SSP offers the best results in terms of balancing the removal of harmful noise and preserving neural information of interest. In cases where only a small number of electrodes are available, an approach based on PCA is deemed helpful. Approaches based on ICA were found to be unsuitable for cardiac artefact removal in ESG, due to a suboptimal balance of artefact removal and signal preservation. Finally, we also approached this issue from a signal-enhancement perspective and observed that in cases where extensive electrode arrays are used in the context of task-based designs, a spatial filtering technique based on canonical correlation analysis (CCA) reveals clear evoked spinal potentials even with single-trial resolution. Taken together, there are several appropriate algorithms for physiological noise removal in ESG, rendering this an accessible and easy-to-use technique for non-invasive assessments of human spinal cord function.

## 1. Introduction

The spinal cord is a key communication pathway between the brain and the peripheral nervous system and is of relevance for sensory, motor and autonomic function (Hochman, 2007). Recent research in animal models has demonstrated that extensive somatosensory processing occurs in the dorsal horn of the spinal cord (Abraira et al., 2017; Paixão et al., 2019), yet direct investigations in humans remain rather scarce due to difficulties in non-invasively imaging this structure. There is, however, a pressing need for such recordings, not only to investigate human spinal cord function in health, but also to understand pathological changes that underlie conditions such as chronic pain, multiple sclerosis and spinal cord injury (Ahuja et al., 2017; Ciccarelli et al., 2019; Kuner and Flor, 2017).

Of the currently available approaches for non-invasive recordings, functional magnetic resonance imaging of the human spinal cord (Kinany et al., 2022; Landelle et al., 2021) offers superior spatial resolution, but is fundamentally limited by its indirect nature (due to neurovascular coupling) and its coarse temporal resolution. These aspects have been addressed using magnetospinography (MSG) based on superconducting quantum interference devices (Adachi et al., 2021; Sumiya et al., 2017). Though successful, these systems are employed in less than a handful of laboratories worldwide, are costly to implement and are not yet commercially available. Alternative MSG recordings based on optically pumped magnetometers (OPMs) have also been proposed (Mardell et al., 2022), yet this technology is currently only in a nascent stage. In contrast to these more recent approaches, there is substantial literature spanning several decades on the non-invasive recording of spinal cord somatosensory evoked potentials (SEPs) using readily-available electrospinography technology (ESG; for reviews, see Cruccu et al., 2008; Mauguière, 2000; Yamada et al., 1980). Such a set-up typically involves transcutaneous electrical stimulation of peripheral nerves and spinal recordings via surface electrodes placed on the neck or the lower back, depending on the responses of interest.

However, in such electrospinographic (ESG) recordings, physiological noise of myogenic and especially cardiac origin is highly detrimental due to the proximity of the recording electrodes to the source of physiological noise, i.e. the heart in the latter case (Cracco, 1973; Cracco et al., 1979; Jones and Small, 1978). In the cervical spinal cord SEPs generally have an amplitude of approximately one to two microvolts (Cruccu et al., 2008), while the cardiac artefact reaches up to several hundred microvolts and has overlapping frequency content with SEPs, illustrating the impact of cardiac noise. Of note is that this detrimental influence is much reduced when electroencephalographic (EEG) recordings are performed on the scalp, thus presenting a challenge unique to ESG recordings (Figure 1).

**Figure 1:**
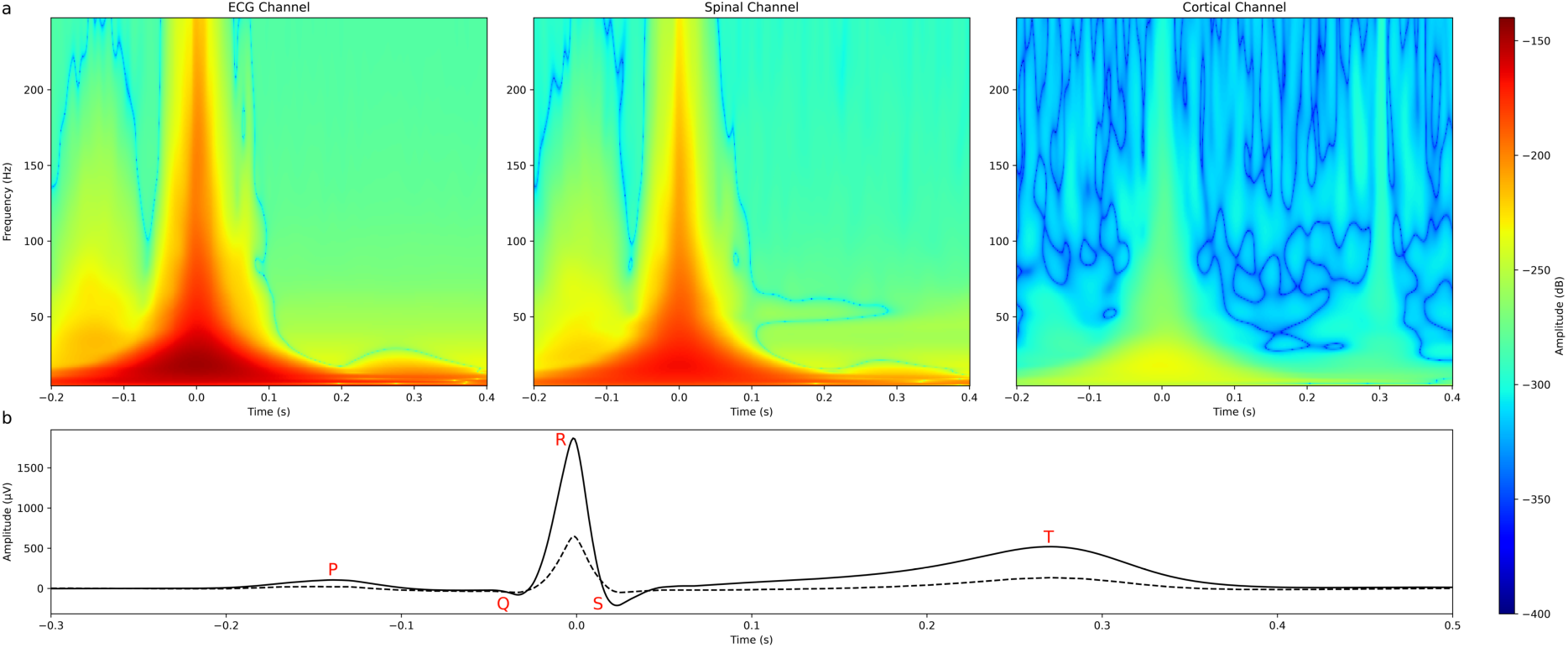
(a) Grand average (N = 36) time-frequency plots demonstrating the manifestation of the heartbeat in the ECG recording channel, with the corresponding effect of the heartbeat in a spinal (L1) and cortical (CPz) channel of interest. The amplitude difference between the spinal and cortical channels shows how detrimental the cardiac artefact is in spinal, as compared to cortical channels. (b) A corresponding time-domain depiction of the cardiac artefact in the ECG channel (solid) and the spinal channel of interest (dashed), with key components of the heartbeat shown in red lettering. In all plots, the R-peak occurs at 0s (note that cortical data are not shown in the time-domain depiction as they would essentially consist of a flat line due to the plot scaling).

The confounding influence of the cardiac artefact is traditionally mitigated in ESG studies via i) employing cardiac-gated stimulation, ii) averaging across large numbers of trials, or iii) extensive high-pass filtering. While effective, each of these approaches has drawbacks: i) cardiac gating does not allow for studies where cardiac-somatosensory interactions are of interest (Al et al., 2020) ii) the necessary number of trials to perform large-scale averaging is unsuitable in modern cognitive neuroscience paradigms (Gijsen et al., 2021) and iii) high-pass filtering is undesirable in the case of resting-state recordings (Mantini et al., 2007). Evidently, there is significant need for an approach that reduces the detrimental impact of the cardiac artefact without such limitations.

There are several approaches known to remove cardiac artefacts in other types of neurophysiological recordings, but to our knowledge, there has been no systematic investigation into the suitability of these methods for non-invasive ESG data. Here, we thus explored the ability of four different algorithms to reduce the impact of the cardiac artefact on ESG data. First, we adapted the principal component analysis optimal basis sets (PCA-OBS) approach pioneered by Niazy et. al. (Niazy et al., 2005), since both the ballistocardiographic (BCG) artefact in simultaneous EEG-fMRI recordings and the cardiac artefact in ESG recordings are signals which can vary in shape at each occurrence but are time-locked to the cardiac cycle. Second, we employed independent component analysis (ICA), which is one of the most popular methods for removing artefacts and has been employed for cardiac artefact removal in both EEG data (Al et al., 2020) as well as invasively-recorded spinal data (Wang et al., 2021). Third, we investigated the potential of signal space projection (SSP: Uusitalo and Ilmoniemi, 1997), a technique that has previously been implemented to remove cardiac artefacts from EEG and ESG data (Haumann et al., 2016; Nierula et al., 2024). Finally, we used canonical correlation analysis (CCA), which can extract high quality evoked potentials in cortical EEG data (Stephani et al., 2020; Waterstraat et al., 2015) - while this method does not directly target the cardiac artefact, it might be possible to obtain reliable SEPs even in the absence of dedicated cardiac artefact correction with this approach. Taken together, while all algorithms have been successfully applied in cortical recordings, their efficacy for cardiac artefact removal in non-invasive surface recordings from the spinal cord remains unknown and is the focus of this investigation.

## 2. Methods

### 2.1. Data acquisition

The data used here come from 36 healthy right-handed volunteers (18 female) who provided written informed consent prior to participation in an experiment, in which they received electrical mixed nerve stimulation of the median nerve at the left wrist and of the tibial nerve at the left ankle (Nierula et al., 2024). Electrical stimulation consisted of a 0.2ms square-wave pulse delivered by constant-current stimulators (DS7A, Digitimer Ltd, Hertfordshire, UK; one stimulator for each nerve) via two bipolar stimulation electrodes to either the median nerve at the left wrist or the tibial nerve at the left ankle. Stimulus intensities were adjusted on an individual level to be just above the motor threshold, but not deemed to be painful. Stimulation was provided with an inter-stimulus interval of 763ms and a jitter of at most +/- 50ms. A total of 2000 stimuli were delivered in alternating blocks of 500 stimuli to either the median or tibial nerve.

The dataset, publicly available at OpenNeuro (https://openneuro.org/datasets/ds004388), includes simultaneously recorded electroencephalography (EEG), electrospinography (ESG), electroneurography (ENG), electromyography (EMG), electrocardiography (ECG) and respiratory information. For the purpose of this study, the focus is on the ESG data with ECG data being used for assessing cardiac activity. For the ESG recordings, two separate patches of 17 electrodes each were placed on a participant’s neck and back: one patch was centred on the cervical spinal cord around an electrode placed over the spinous process of the 6^th^ cervical vertebra and another patch was centred on the lumbar spinal cord around an electrode placed over the spinous process of the 1^st^ lumbar vertebra. The electrodes were arranged along the midline of the vertebral column and extended laterally from there, with the ESG reference electrode being located between the cervical and lumbar spinal electrodes over the spinal process of the 6^th^ thoracic vertebra; two ventral electrodes were also placed to enable anterior re-referencing (more details on patch construction are available in the original publication: Nierula et al., 2024). For median nerve stimulation, channels of interest were determined to be SC6 (the central cervical electrode located over the spinous process of the 6th cervical vertebra) as well as S6 and S14 (the immediately neighbouring electrodes directly above and below SC6 along the vertebral column). For tibial nerve stimulation, channels of interest were determined to be L1 (the central lumbar electrode located over the spinous process of the 1st lumbar vertebra) as well as S23 and S31 (the immediately neighbouring electrodes directly above and below L1 along the vertebral column). These channels represent midline channels in the array, and generally have the strongest somatosensory evoked potentials due to their positioning.

### 2.2. ESG preprocessing

Unless mentioned otherwise, all analyses were performed using Python 3.9 and MNE (https://mne.tools/stable/index.html; version 1.0.3), an open-source toolbox for analysing human neurophysiological data (Gramfort, 2013). First, the data was down-sampled to 1000Hz. Next, in order to remove artefacts resulting from electrical stimulation, data were linearly interpolated between −7ms and 7ms relative to stimulus onset. Data after this stage of processing is referred to as ‘Uncleaned’ from now on. Both this Uncleaned data, and the data after each method of cardiac artefact removal were notch filtered around 50Hz with an FIR zero-phase filter and bandpass filtered between 1Hz and 400Hz with a 2^nd^ order Butterworth zero-phase filter. A graphical overview of the processing and analysis steps – including the employed correction methods as well as the metrics used for their evaluation – is provided in Figure 2.

**Figure 2:**
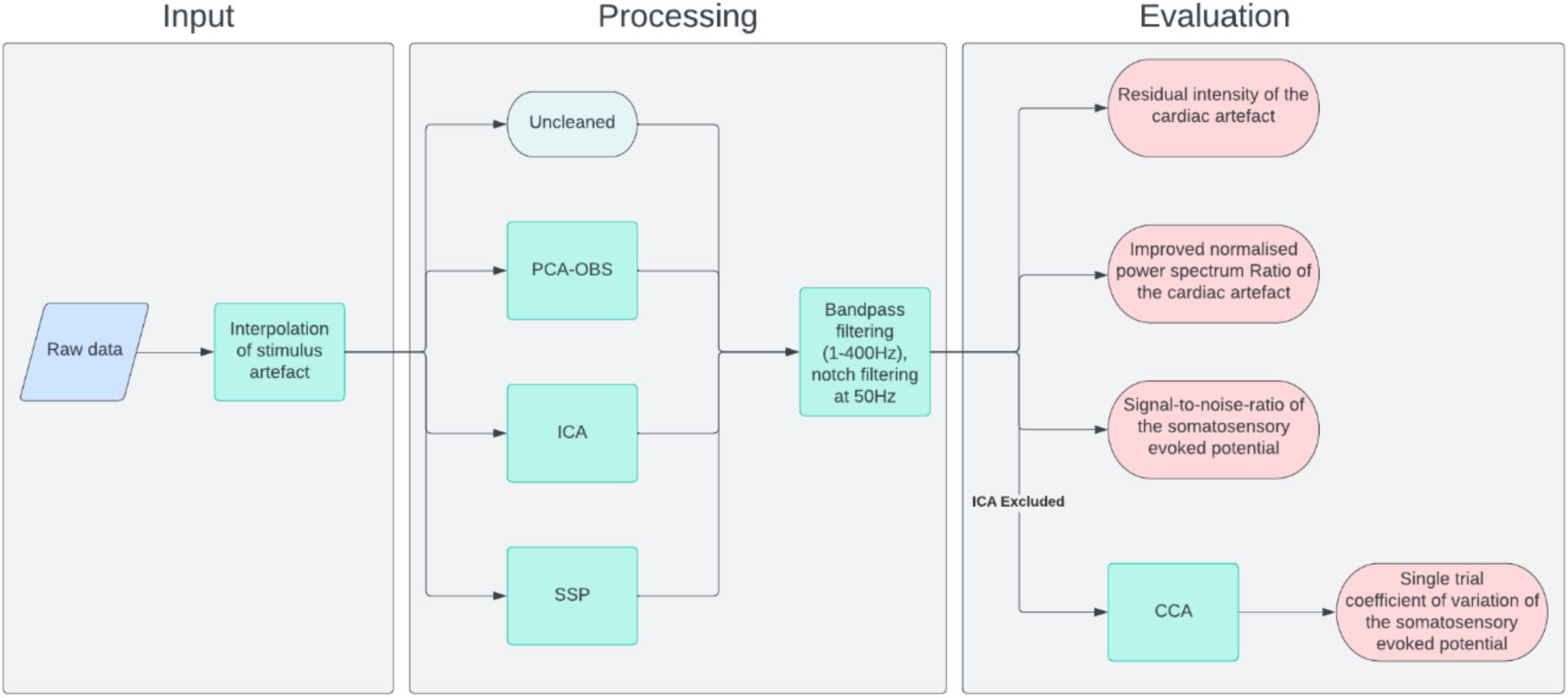
Overview of methods implementation. First (left panel), the input data is pre-processed to remove the stimulation artefact. Data after this stage is referred to as Uncleaned. This Uncleaned data then (middle panel) has the cardiac artefact removed by either principal component analysis optimal basis sets (PCA-OBS), independent component analysis (ICA) or signal space projection (SSP). In each case, the data is then bandpass filtered between 1Hz and 400Hz, and notch filtered at 50Hz to remove powerline noise. Each algorithm is then (right panel) evaluated based on its ability to remove the cardiac artefact (using the residual intensity and the improved normalised power spectrum ratio) and its ability to preserve the somatosensory evoked potentials of interest (using the signal-to-noise ratio). In a final step, the ability of canonical correlation analysis (CCA) to create meaningful somatosensory evoked potentials either with or without pre-cleaning of the cardiac artefact is tested. To achieve this aim, CCA is run on the Uncleaned data, as well as the PCA-OBS and SSP cleaned data. The performance of CCA is then quantified by means of the single trial coefficient of variation of the somatosensory evoked potentials which describes the variation in single-trial amplitudes across trials.

### 2.3. Cardiac artefact removal techniques

#### 2.3.1. Principal Component Analysis Optimal Basis Sets (PCA-OBS)

This method is an adaptation of the method pioneered by Niazy and colleagues (Niazy et al., 2005) for the removal of the ballistocardiographic artefact arising from cardiac activity in simultaneous EEG-fMRI recordings. It works on each channel separately and assumes each cardiac artefact occurrence in each EEG channel is independent of all previous occurrences in time. Principal component analysis is performed on a matrix of all artefact occurrences to capture the principal variations of the artefact. Then, the first *n* principal components can be selected as an optimal basis set, which can be fit to each artefact occurrence using Piecewise Cubic Hermite Interpolating Polynomials (PCHIP) and subtracted from the signal.

While the original implementation for simultaneous EEG-fMRI is available as a plug-in for EEGLAB (https://fsl.fmrib.ox.ac.uk/eeglab/fmribplugin/), we implemented this method via custom-written scripts in Python 3.9 to use it as a correction method for ESG. The timing of each R-peak was previously determined using automatic detection and manual correction for each participant (Nierula et al., 2024), and these latencies were used again here. A window around each R-peak was selected based on the median RR interval (the time between successive R-peaks) for each participant and determines the area in which the fitted artefact is created for each heartbeat occurrence. Following the original implementation, the data was then high pass filtered at 0.9Hz before PCA was performed – afterwards the first four components identified by PCA as well as the mean effect were selected to form the optimal basis set. The original Niazy algorithm (Niazy et al., 2005) removes 3 principal components to remove the ballistocardiographic artefact, however in our case, previous analyses (Nierula et al., 2024) revealed removing 4 principal components works well for cardiac artefact cleaning in our dataset. This optimal basis set is then fit to each occurrence of the artefact to form a fitted artefact, which is directly subtracted from the ESG data in each channel.

Due to the subtraction of the fitted artefact, there can be sharp edges at the beginning and end of fitting windows. To counteract this effect, the PCA-OBS algorithm was further modified here by multiplying the fitted artefact in each window with a Tukey window (α = 0.25) prior to subtraction to investigate potential reduction of these edge effects.

#### 2.3.2. Independent Component Analysis (ICA)

ICA is a blind source separation technique for transforming an observed multidimensional random vector into components that are statistically as independent from each other as possible (Hyvarinen, 1999). ICA was applied to the Uncleaned data using the Fast-ICA method (Hyvarinen, 1999), with the number of components specified as the number of ESG channels present across both spinal patches.

The components to be removed were automatically determined in MNE, which selects components for removal based on cross-trial phase statistics. Cross trial phase distributions are time-dependent histograms, and, in this case, these are phase histograms calculated across trials, where each trial is defined by a 1 second time-window centred on the R-peak of the independent components. Independent components associated with cardiac activity will be synchronous with the ECG trace and therefore exhibit a non-uniform cross-trial phase distribution (Dammers et al., 2008). The independent components are then selected for removal based on the significance value of the Kuiper statistic and the data is finally reconstructed by zeroing the excluded components and inverse transforming the data. In cortical data, this approach has been shown to be robust and highly sensitive for the removal of components attributed to cardiac activity as determined from the ECG channel (Dammers et al., 2008).

Due to the poor performance of ICA, alternative ICA procedures were also tested: i) applying ICA to anteriorly re-referenced data and ii) applying ICA separately to each spinal patch (Supplemental Material). Further brief attempts to use ICA for cardiac artefact removal were made by considering a hyperbolic tangent function to perform ICA, and by performing ICA separately depending on the stage of respiration. These algorithms were not systematically tested due to the lack of success encountered with test participants in this dataset, and as such they were not pursued further and are not presented in the context of this work.

#### 2.3.3. Signal Space Projection (SSP)

SSP (Uusitalo and Ilmoniemi, 1997) assumes the signals produced by different sources have different and fixed orientations in sensor space, which implies each source has a distinct and stable field pattern. Since SSP is a spatial filtering technique, the activity from both spinal patches are considered in tandem to maximise the spatial information available.

Measured signals can be divided into an estimate of the signals produced by a distinct source (here: cardiac activity) and all other sources, including spinal signals of interest. For cardiac artefact removal, SSP was performed on epochs created from 0.2s before to 0.4s after the identified R-peaks, thus allowing the identification of projection vectors associated with the sources of cardiac activity which can subsequently be removed.

An exploratory analysis was conducted to determine the number of projectors to remove, with all numbers between 1 and 20 investigated – 20 was chosen as an upper limit, as we noticed that the signal-to-noise ratio of the SEPs of interest had decreased substantially. Ultimately, six projectors were selected for removal (with this approach simply referred to as SSP from now on) based on three metrics; please see Supplementary Material for further details (Table S1; Table S2).

### 2.4. Canonical Correlation Analysis (CCA)

All previous approaches (PCA-OBS, ICA, SSP) are specifically aimed at removing cardiac artefacts and thus are equally suitable for both resting-state and task-based recordings. However, in the context of task-based recordings as employed here, it is also possible to focus on enhancing the signal of interest, rather than removing data associated with detrimental noise. This can be achieved using spatial filtering approaches such as CCA (Fedele et al., 2013) which has been successfully applied both cortically (Stephani et al., 2020; Waterstraat et al., 2015) and spinally (Nierula et al., 2024) to enhance signals of interest.

CCA was performed to extract single-trial SEPs using the Modular EEG Toolkit (MEET; https://github.com/neurophysics/meet; Waterstraat et al., 2015) and is used to find the weights *w_x_* and *w_y_* that mutually maximise the correlation

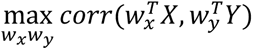

where X and Y are both multi-channel signals. We used a variant of CCA also known as canonical correlation average regression (Waterstraat et al., 2015) – in this case, X is a two-dimensional matrix that contains all single-trial epochs from 1 to N for each channel, and Y contains N times the average SEP for each channel. With this construction, the weight matrix *w_x_* represents the spatial filters that in combination with *w_y_* maximise the correlation between the single trial activity (X) and the average SEP (Y). Taking into account knowledge of the expected latency of SEP’s evoked by median (13ms) and tibial (22ms) nerve stimulation, the spatial filters *w_x_* were trained using short segments after stimulus delivery (median nerve stimulation: 7-37ms, tibial nerve stimulation: 7-47ms; these time windows were chosen in order to have ~25ms of data after the expected spinal potential in order to have sufficient data for an accurate assessment of the noise covariance), but the resulting spatial filters were applied to whole length of the epochs. Further, for each type of peripheral nerve stimulation, only electrodes from the relevant spinal patch were included (median nerve: cervical patch; tibial nerve: lumbar patch). As CCA is insensitive to the polarity of the signal, if the potential at the expected latency of the time course of the chosen component manifested as a positive peak, the data was inverted.

We applied CCA separately in the lumbar and cervical spinal patches, for i) Uncleaned, ii) PCA-OBS-cleaned and iii) SSP-cleaned data to investigate whether CCA can provide additional benefits when combined with a dedicated artefact correction method, or whether it is sufficient by itself (note that ICA-cleaned data were not used here due to the poor performance of this method). A single CCA component from the top four CCA components (ranked by their canonical correlation coefficient) was selected for each participant and each method. To select the optimal component, three criteria we considered; i) there must be a peak at the expected SEP latency in the component’s average time-course, ii) consistent peaks at the expected latency are identifiable in individual trials, and iii) the spatial pattern (computed by multiplying the covariance matrix of X by the spatial filters w_x_, to take the data’s noise structure into account (Haufe et al., 2014)) should match the expected spatial topography (irrespective of polarity) for cervical or lumbar spinal cord SEPs, respectively.

### 2.5. Metrics

The performance of PCA-OBS, ICA and SSP was quantified in two different ways. On the one hand, we assessed each method’s efficacy of cardiac artefact reduction in two ways, first, in the time domain via the remaining amplitude of the cardiac artefact, referred to by the metric residual intensity (Marino et al., 2018) and second, in the frequency domain using the improved normalised power spectrum ratio of the cardiac artefact (Javed et al., 2017). On the other hand, we assessed each method’s ability to preserve or enhance the response of interest by computing the signal-to-noise ratio of the SEPs. Furthermore, the effect of CCA was examined in terms of the SEPs’ amplitude variation across trials in the potential window of interest using the coefficient of variation.

All these metrics were computed at the level of individual participants; group-level results are reported as the average ± the standard error of the mean computed across participants. Apart from the last metric (coefficient of variation after CCA cleaning), all metrics were investigated in channels S6, SC6 and S14 for median nerve stimulation and channels S23, L1 and S31 for tibial nerve stimulation.

#### 2.5.1. Residual Intensity (RI)

The data of the channels of interest was epoched from −300ms to 400ms with respect to the detected R-peaks in order to not only include the QRS-complex, but also the P- and T-waves, and baseline corrected within a period of −300ms to −200ms (this baseline period was chosen as it should occur before the P-wave in most participants, without risking overlap with the T-wave of the preceding heartbeat). All epochs were then averaged for each channel to form the residual evoked response. For each method (PCA-OBS, ICA, SSP) the residual intensity was computed for each channel by taking the root mean square (RMS) of the cleaned data and dividing it by the RMS of the Uncleaned data and then multiplying by 100 to form a percentage (with smaller values indicating a better performance).

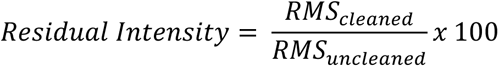

Last, the residual intensity computed for the three channels of interest for each electrode patch was averaged to form the final metric.

#### 2.5.2. Improved Normalised Power Spectrum Ratio (INPSR)

For each participant, the fundamental frequency of the heartbeat was computed, along with the frequency of the first four harmonics. The power in a 0.2Hz band centred on the fundamental frequency and its harmonics was then computed using the Yet Another Spindle Algorithm (YASA; Vallat and Walker 2021) for each channel and each method (PCA-OBS, ICA, SSP). The power within these predefined spectral ranges was then summed to obtain the total band power per channel and method. The INPSR was then computed for each by dividing the total band power of the Uncleaned data by the total band power of the cleaned data (with larger values indicating better performance).

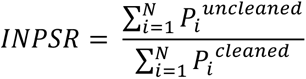

Last, the INPSR computed for the three channels of interest for each electrode patch was averaged to form the final metric.

#### 2.5.3. Signal-to-Noise Ratio (SNR) of SEPs

The data was epoched from −200ms to 700ms around each stimulus onset with a baseline period from −100ms to −10ms relative to stimulus onset. These epochs were averaged across all trials to form the evoked response in each channel of interest. We then defined time-windows of interest around the typical response latency (5ms to either side of the typical N13 response and 10ms to either side of the typical N22 response; note that the larger time window for the N22 is chosen due to the higher latency variability induced by tibial stimulation) and searched for the strongest negative deflection within these time-windows. The magnitude of the largest negative deflection across the channels of interest was chosen to be the peak amplitude. The standard deviation of the baseline period was then computed for the relevant channel, and the SNR was taken to be the peak amplitude divided by the standard deviation in the baseline period.

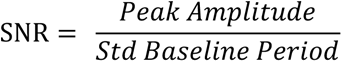

#### 2.5.4. Coefficient of Variation (CoV)

For each dataset that CCA was applied to (Uncleaned, PCA-OBS, SSP), the epochs returned after CCA were cropped according to the type of stimulation (8-18ms for median nerve stimulation and 12-32ms for tibial nerve stimulation). The peak-to-peak amplitude taken as the difference between the maximum negativity and the maximum positivity within each trial was then computed, and the CoV was calculated based on the variation of the peak-to-peak amplitude across all trials. The CoV across trials was computed for each method both before and after the application of CCA in order to compare the impact of pre-cleaning the cardiac artefact prior to the application of CCA.

### 2.6. Statistics

Considering that this dataset was acquired with relatively regular electrical stimulation (an inter-stimulus interval of 763ms and a jitter of at most +/-50ms), we first assessed whether the cardiac cycle was locked to our electrical stimulation. Here, a Rayleigh test was used to determine whether or not there is significant (p<0.05) locking of the cardiac cycle to the tibial or median nerve stimulation, using the angle from the current stimulus to the last occurring R-peak (performed using pycircstat: https://github.com/circstat/pycircstat).

Whenever statistical significance (p<0.05) was assessed with respect to the calculated metrics (SNR, RI, INSPR, CoV), non-parametric permutation testing was performed (with the number of permutations set to 2000). One-sample, two-tailed permutation t-tests were performed separately on each metric (RI, INPSR and SNR) to compare the cardiac artefact denoising methods’ performance. Pairings were formed by subtracting each participant’s metric after cardiac artefact denoising (PCA-OBS, ICA or SSP) from that participant’s metric prior to cleaning or after a different method of cleaning. The max-T method was used to adjust the p-values to account for the number of tests within each metric, i.e. controlling the family-wise error rate (Nichols and Holmes, 2002). The same statistical procedure was performed when investigating the effect of CCA on the coefficient of variation across trials.

## 3. Results

### 3.1. Cardiac cycle locking to stimulation

When the angle from the stimulus to the previous heartbeat is considered for all trials and participants, neither the median nerve (p = 0.55) nor the tibial nerve condition (p = 0.34) showed a significant result. When testing each participant (N = 36) and condition (median nerve, tibial nerve) separately, only three of the 72 conditions were significant (p_uncorrected_ < 0.05). Overall, we can thus be confident that there was minimal locking of the cardiac cycle to the somatosensory stimulation.

### 3.2. Quantification of SEP and cardiac artefact amplitudes

While the cardiac artefact in both the cervical and lumbar spinal cord is known to be substantially larger than the SEPs of interest, to our knowledge no quantitative data exist on this yet and we thus set out to determine the respective amplitudes in our specific dataset. For this, we used the central electrode of the cervical and lumbar electrode patches (SC6 and L1, respectively) of the Uncleaned data. The average amplitude of the cardiac artefact in the cervical spinal cord was −297µV, while the cervical N13 response had an average amplitude of −0.86µV. Similarly, in the lumbar spinal cord the average amplitude of the cardiac artefact was 657µV while the lumbar N22 response had an average amplitude of −0.80µV. Thus, on average the cardiac artefact is 345 times larger than the SEPs of interest in the cervical spinal cord, and 819 times larger in the lumbar spinal cord.

### 3.3. Cardiac artefact reduction

First, we qualitatively investigated the reduction of the cardiac artefact via PCA-OBS, ICA and SSP at the group-level in the time domain (Figure 3) as well as in the frequency domain (Figure 4). Each method achieved a marked decrease in the amplitude and power associated with the cardiac artefact, with ICA visually offering the best performance in the cervical spinal channel of interest (SC6), while SSP appears to offer superior performance in the lumbar spinal channel of interest (L1). Despite these differences, each method was individually successful, with a reduction in the peak magnitude of the R-peak in the grand-averaged waveform in cervical spinal channels from ~300µV to less than 0.5µV, while in the lumbar spinal cord there is a reduction from ~650µV to less than 3µV in all cases, i.e. a ~500 fold and ~200 fold reduction in artefact amplitude at the group level for cervical and lumbar spinal cord data, respectively (exemplary single-participant time-courses can be found in the Supplementary Material). These time-domain results (Figure 3) are supported by the time-frequency representations (Figure 4), which show a severely reduced power associated at frequencies and time points known to correspond to cardiac activity (as seen in Figure 1). Further, edge-effects induced by PCA-OBS are visible, with sharply increasing signal amplitude observable at the border of the artefact window at 0.4s (an investigation aimed at addressing these edge-effect via Tukey Windows can be found in the Supplementary Material).

**Figure 3:**
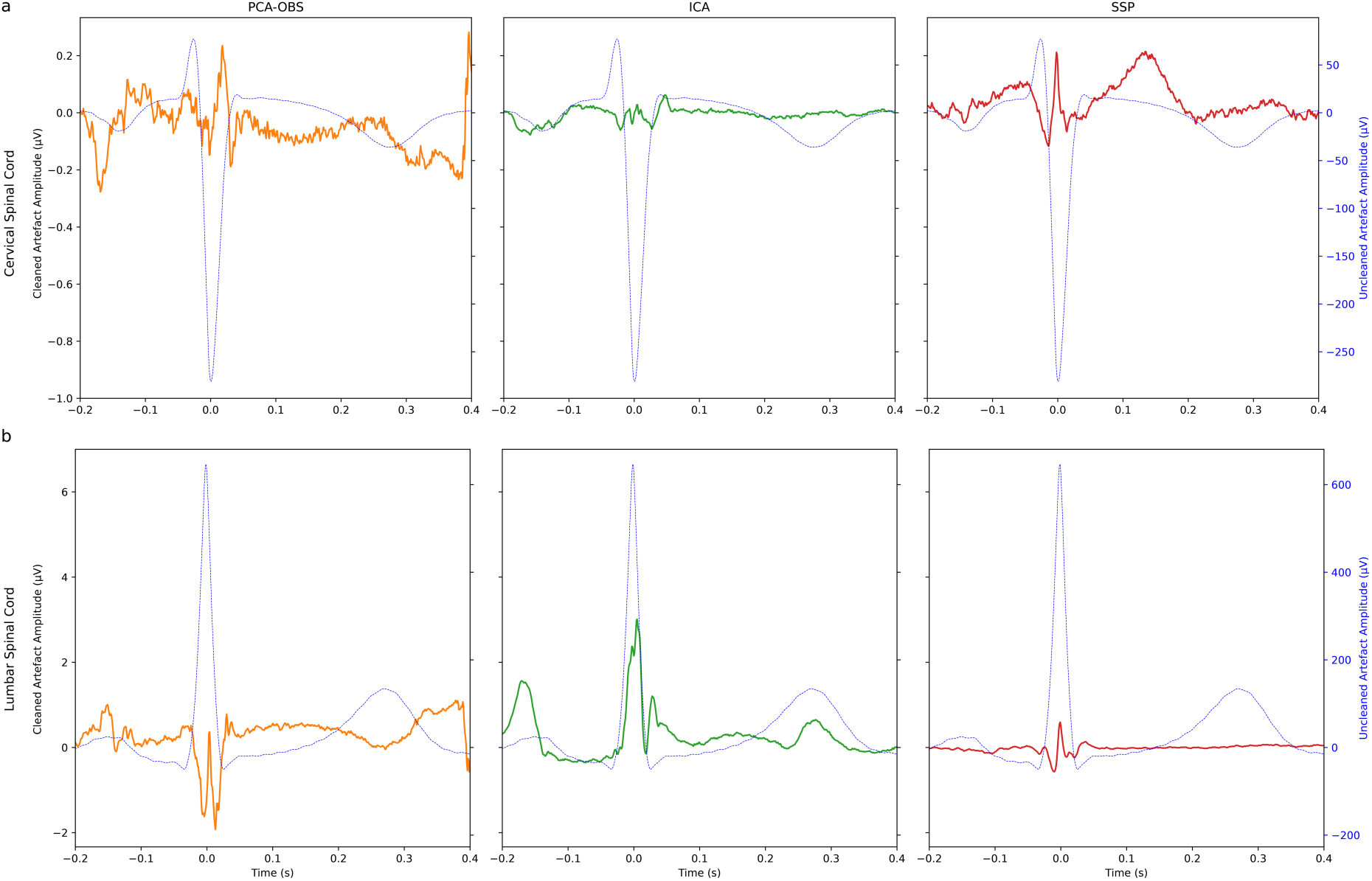
Cardiac artefact before and after cleaning. The group-level cardiac artefact after each method of cardiac artefact correction in the cervical (channel SC6; a) and the lumbar (channel L1; b) spinal cord, with the time course of the Uncleaned artefact shown in blue. The scale bar on the left of each row refers to the coloured traces depicting the cardiac artefact after cleaning, while the scale bar on the right of each row refers to the Uncleaned cardiac artefact (blue). Please note the vastly different scale between cleaned and Uncleaned data, as well as the difference in the amplitudes in the cervical and lumbar spinal cord. 0ms corresponds to the centre of the R-peak of the heartbeat.

**Figure 4:**
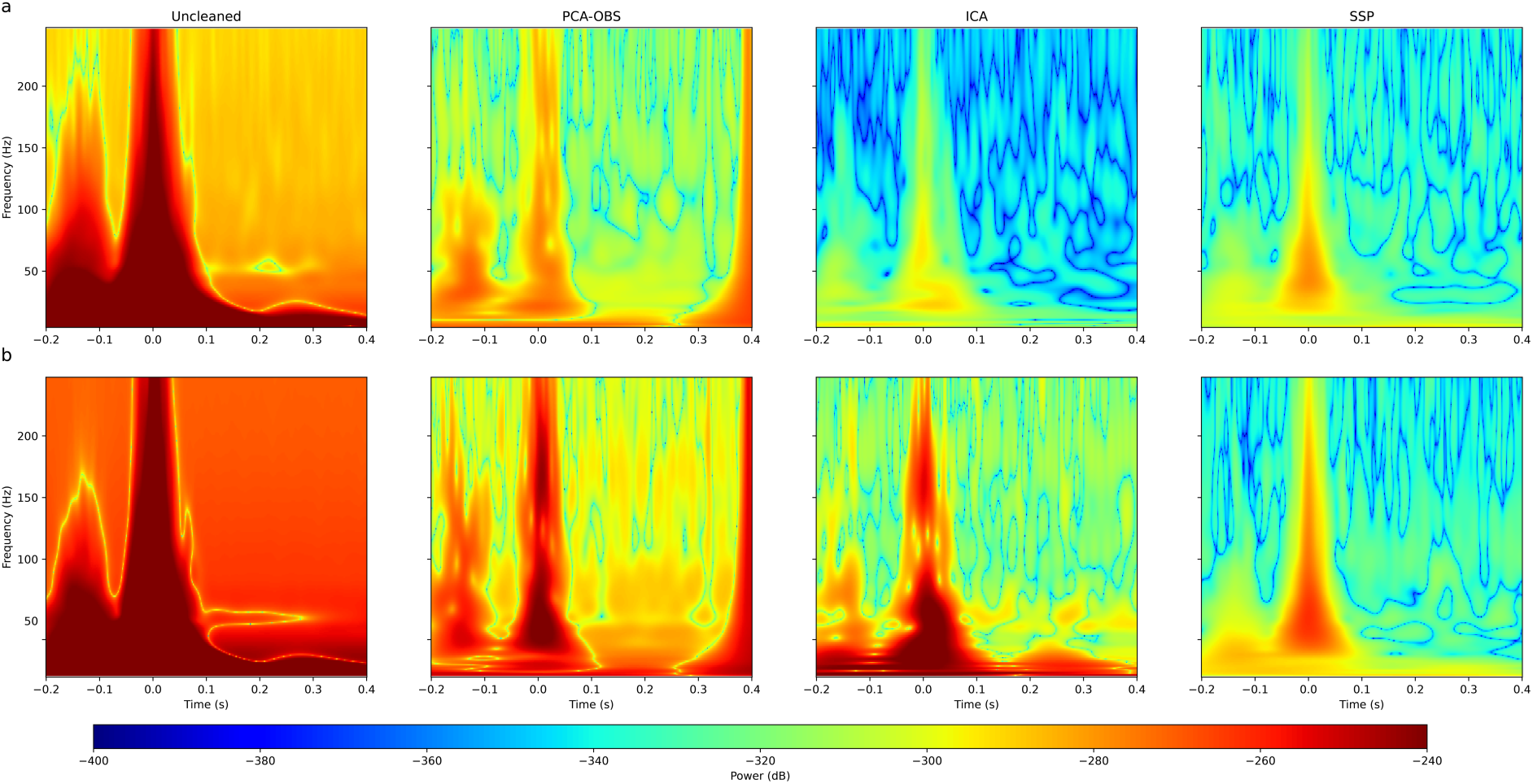
Time-Frequency representations of the grand average cardiac artefact across all participants and all trials both before and after each method of cardiac artefact cleaning in the cervical spinal cord (channel SC6; a) and lumbar spinal cord (channel L1; b). 0ms corresponds to the centre of the R-peak of the heartbeat.

In order to support these descriptive results, we next turned to formally assessing each method via artefact-related metrics, namely the residual intensity (RI) and the improved normalised power spectrum ratio (INPSR) of the cardiac artefact (Table 1). The RI results indicate that each method is able to reduce the artefact to less than 1% (except for PCA-OBS in lumbar data, which was ~1.2%), with ICA offering the best performance in cervical data (followed by SSP) and SSP offering the best performance in lumbar data. In the cervical spinal cord, the difference between ICA and PCA-OBS is found to be significant (p = 0.003) alongside the difference between ICA and SSP (p = 0.001), while the difference between SSP and PCA-OBS is not significant (p = 0.18). In the lumbar spinal cord, the reverse is true: the difference in the performance of ICA versus PCA-OBS is not significant (p = 0.90) alongside the difference between ICA and SSP (p = 1.0), while the difference between SSP and PCA-OBS is significant (p < 0.001).

**Table 1:** Group-level residual intensity (RI), improved normalised power spectrum ratio (INPSR) and SEP signal-to-noise ratio (SNR) results for cervical and lumbar channels of interest in response to median and tibial nerve stimulation, respectively. The standard error of the mean (across participants) is also reported.

| Cervical Spinal Cord |  |  |  |
| --- | --- | --- | --- |
|  | RI | INPSR | SNR |
| Uncleaned | - | - | $2.45 \pm 0.33$ |
| PCA-OBS | $0.69 \pm 0.14\%$ | $4.55\text{E}+02 \pm 6.33\text{E}+01$ | $7.67 \pm 0.83$ |
| ICA | $0.19 \pm 0.06\%$ | $2.95\text{E}+90 \pm 2.88\text{E}+90$ | $3.30 \pm 0.54$ |
| SSP | $0.43 \pm 0.04\%$ | $3.81\text{E}+03 \pm 9.14\text{E}+02$ | $14.30 \pm 1.27$ |
| Lumbar Spinal Cord |  |  |  |
|  | RI | INPSR | SNR |
| Uncleaned | - | - | $1.70 \pm 0.23$ |
| PCA-OBS | $1.16 \pm 0.30\%$ | $6.36\text{E}+02 \pm 7.97\text{E}+01$ | $4.65 \pm 0.56$ |
| ICA | $0.58 \pm 0.53\%$ | $8.51\text{E}+93 \pm 8.51\text{E}+93$ | $2.09 \pm 0.23$ |
| SSP | $0.29 \pm 0.02\%$ | $2.90\text{E}+04 \pm 4.68\text{E}+03$ | $9.80 \pm 1.29$ |

The INPSR results are largely concurrent with the RI findings, with the cardiac artefact being at least 455 times smaller after cardiac artefact cleaning, though in this case ICA offers the best performance in both the cervical and lumbar spinal cord, both times followed by SSP. In the cervical spinal cord, the difference between SSP and PCA-OBS is found to be significant (p < 0.001), while the difference between PCA-OBS and ICA is not significant (p = 0.36) alongside the difference between SSP and ICA (p = 0.36). In the lumbar spinal cord, the same is true: the difference between the performance of PCA-OBS and SSP is significant (p < 0.001), while the difference between PCA-OBS and ICA is not significant (p=0.50) alongside the difference between SSP and ICA (p=0.50). While the INPSR associated with ICA appears to be superior, ICA fails to reach significance when compared to both PCA-OBS and SSP in the cervical and lumbar spinal cord – this is due to ICA’s large standard error in the INPSR results.

### 3.4. Somatosensory evoked potentials

While the ability of each method to reduce the impact of the cardiac artefact is of central importance to this work, it is also necessary to determine whether each method can maintain meaningful neural content, i.e. SEPs in our case. Thus, the impact of each method on the SEPs was visually inspected and then quantified by means of the signal-to-noise ratio (SNR) at the level of individual participants.

In the cervical cord, the grand-average time-course demonstrates that even in the absence of cleaning of the cardiac artefact, it is possible to identify the N13 after median nerve stimulation (Figure 5). In the lumbar cord however, it is much more difficult to identify the N22 after tibial nerve stimulation (Figure 6) in the Uncleaned data, where only a small negative deflection is visible. In both instances, the SEPs of interest are largely obscured by low-frequency content in the Uncleaned data, however by employing PCA-OBS and SSP, it is possible to vastly reduce low-frequency activity associated with cardiac activity, thus revealing much clearer somatosensory evoked potentials (see also Supplementary Material).

**Figure 5:**
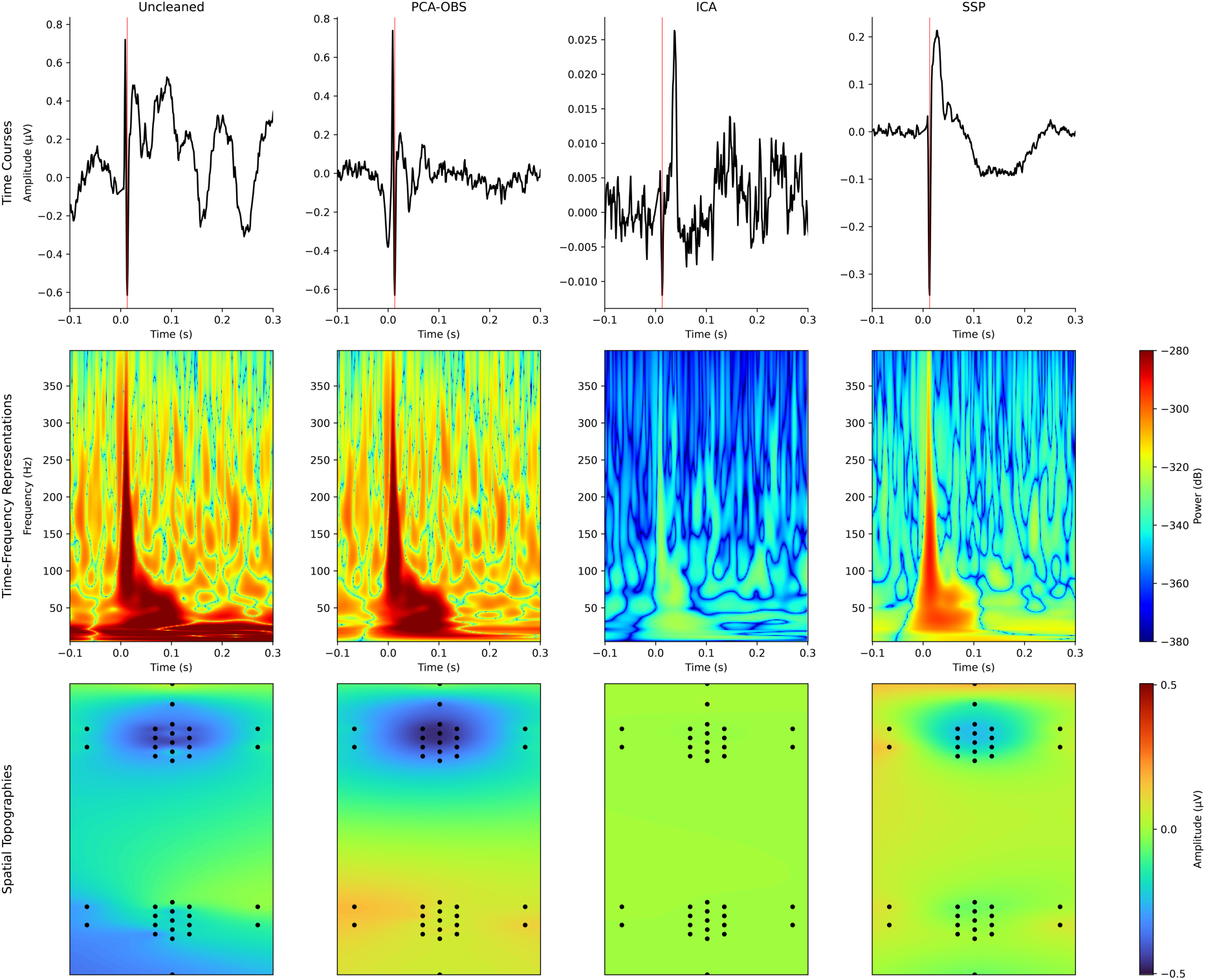
Time courses, time frequency representations and spatial topographies for Uncleaned, PCA-OBS, ICA and SSP corrected data in response to median nerve stimulation (time-courses and time-frequency representations originate from electrode SC6 in the cervical spinal cord). The PCA-OBS data used to generate the time-frequency representation has been linearly interpolated from −7ms to 7ms to avoid artefactual components of no interest obscuring the SEP. The expected SEP latency (13ms) is marked by a red line in the time courses (left). The spatial topography represents the activity from 1ms before to 2ms after the expected peak.

**Figure 6:**
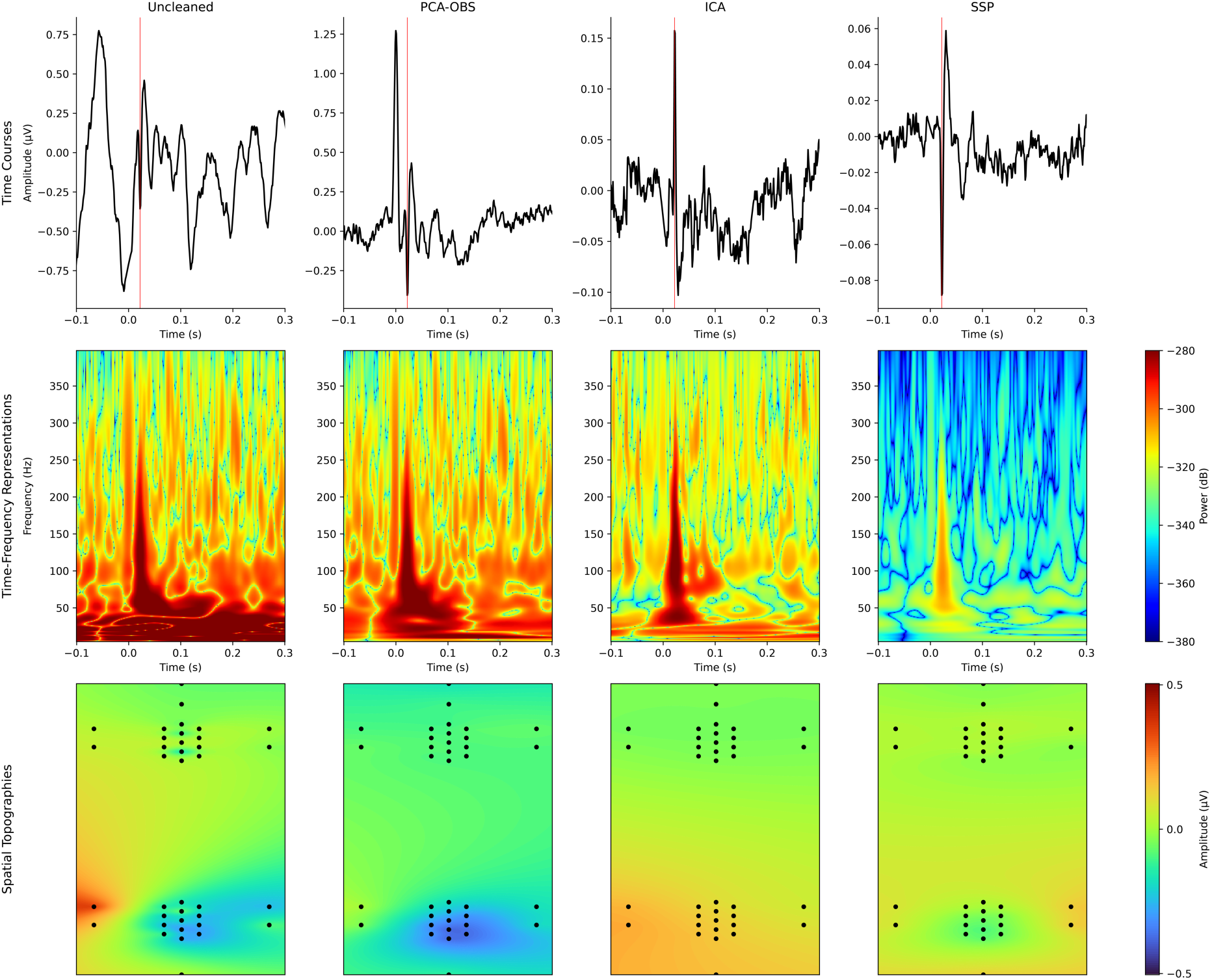
Time courses, time frequency representations and spatial topographies for Uncleaned, PCA-OBS, ICA and SSP corrected data in response to tibial nerve stimulation (time-courses and time-frequency representations originate from electrode L1 in the lumbar spinal cord). The PCA-OBS data used to generate the time-frequency representation has been linearly interpolated from −7ms to 7ms to avoid artefactual components of no interest obscuring the SEP. The expected SEP latency (22ms) is marked by a red line in the time courses (left). The spatial topography represents the activity from 1ms before the expected peak to 2ms after. Channel S34 has been excluded as a bad channel for participant 34.

This effect is further supported by the time-frequency representations (Figure 5: cervical spinal cord; Figure 6: lumbar spinal cord) which in the case of both PCA-OBS and SSP show far lower power at the most dominant frequencies associated with the heartbeat (<30Hz), but maintain clear power peaks at the expected latency of the SEP, while the Uncleaned data shows the clear presence of high power, low frequency noise. Finally, the robustness of each method can be examined by its ability to localise the spinal activity to the correct electrode patch (cervical spinal cord for median nerve stimulation (Figure 5) and lumbar spinal cord for tibial nerve stimulation (Figure 6). This is clearly achieved by both PCA-OBS and SSP, but not by ICA, which also did not allow for a clear identification of the SEPs with the correct polarity in the time-domain and time-frequency representations. This pattern of results is a clear change from the results in reducing the cardiac artefact – and shows that to achieve such convincing reductions of the cardiac activity present in the signal, ICA leaves very little spinal activity remaining. Further testing was conducted to investigate the suitability of alternative ICA arrangements: i) by performing ICA on anteriorly re-referenced data and ii) by including only the spinal patch of interest, but neither had an appreciable effect on the results (Supplementary Material).

Overall, all methods show an improved SNR of the signal of interest (Table 1), with ICA performing worst. The visual impressions from Figure 5 and Figure 6 are supported by the formal SNR results shown in Figure 7 and Table 2, with SSP exhibiting the best performance in both the cervical and lumbar spinal cord (> 5-fold increase in SNR compared to Uncleaned data and nearly 2-fold increase in SNR compared to PCA-OBS; exemplary single-participant time courses can be found in the Supplementary Material). Despite the overall excellent performance of SSP in eliciting high-fidelity SEPs, the absence of the P11 potential from the cervical SEP time-course (Figure 5) is worthy of mention as this is a documented spinal potential after median nerve stimulation (Desmedt and Cheron, 1982). While the reason for this absence is unclear, we note that the time-frequency and spatial topography as well as the time-course of the main cervical SEP component (N13) are as expected. Additionally, a prominence about the stimulation point (0s) in the lumbar spinal cord can be observed in the case of PCA-OBS, which is due to an interaction between interpolation around stimulus onset (to remove the stimulus artefact) and the PCA-OBS fitting algorithm.

**Figure 7:**
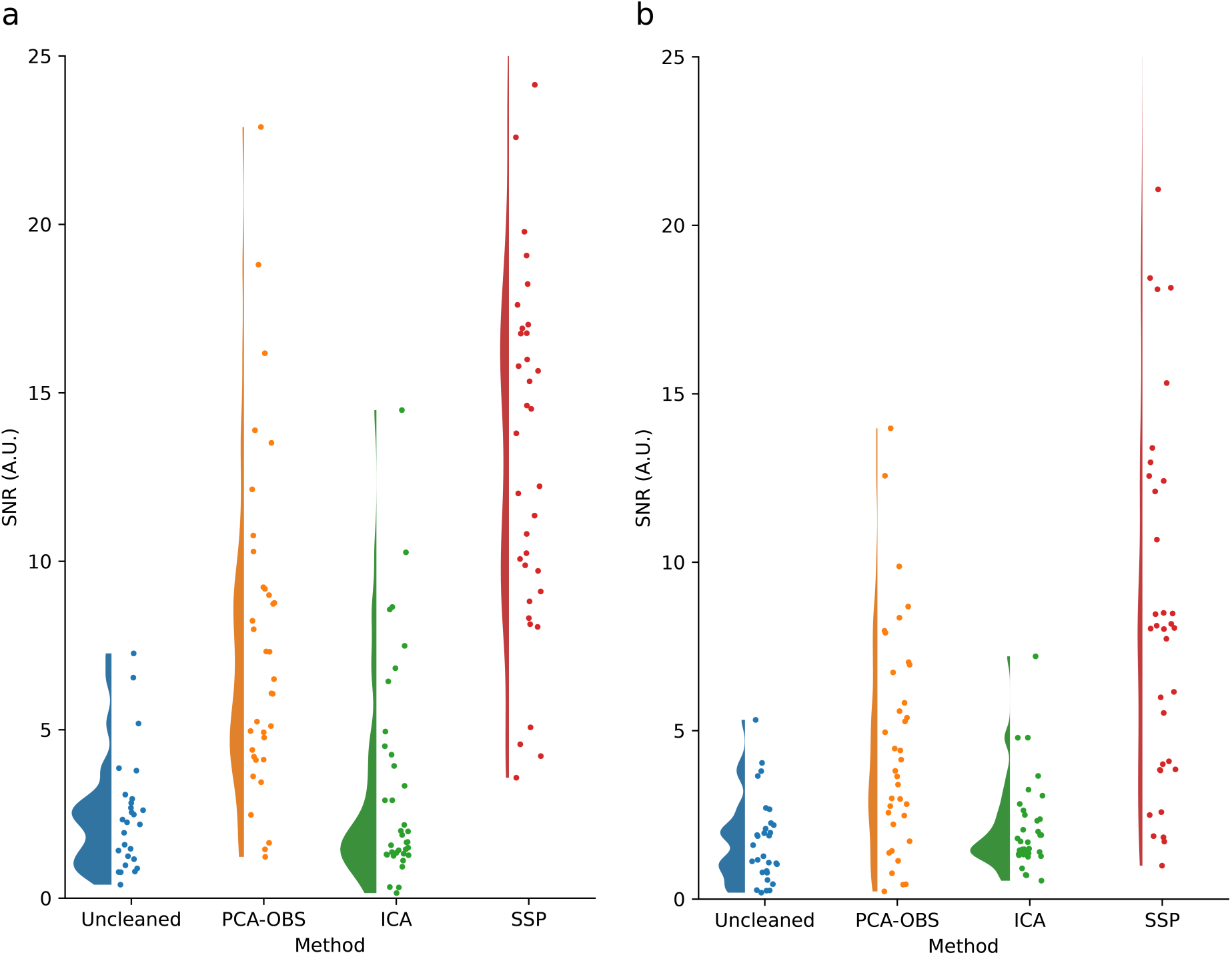
SEP Signal-to-noise ratio (SNR) results for Uncleaned data and each method of cardiac artefact removal in the cervical spinal cord (a) and the lumbar spinal cord (b), with circles indicating participants.

**Table 2:** Statistical comparison of SEP-SNR after different cleaning methods. Reported are two-tailed p-values associated with the difference between the SNR of each pair of methods; significant results are shaded green, while those that do not reach significance are shaded yellow (for underlying SNR values, see Table 1).

|  | Cervical Spinal Cord | Lumbar Spinal Cord |
| --- | --- | --- |
| Uncleaned – PCA-OBS | 0.001 | 0.001 |
| Uncleaned - ICA | 0.490 | 0.210 |
| Uncleaned – SSP | 0.001 | 0.001 |
| PCA-OBS - ICA | 0.001 | 0.006 |
| PCA-OBS – SSP | 0.001 | 0.001 |
| ICA – SSP | 0.001 | 0.001 |

### 3.5. Canonical Correlation Analysis

Having established that SSP offers the best performance - by balancing the removal of cardiac activity with the maintenance of relevant spinal signals - the spatial filtering technique CCA was employed to determine whether dedicated cleaning of the cardiac artefact is even necessary to extract high-quality SEPs in the context of task-based experiments. To quantify this, we determined the coefficient of variation (CoV) across all single trials within each participant which allowed us to determine the sensitivity to detecting SEPs at the single trial level, both before and after the application of CCA. At this point of the analysis, it was decided not to perform CCA on the data where the cardiac artefact was pre-cleaned using ICA, as ICA failed to maintain sufficient spinal activity of interest (see above). Thus, the results presented in this section include only conditions were CCA was applied to Uncleaned, PCA-OBS cleaned and SSP cleaned data.

CCA is seen to clearly reduce the single-trial variation, as in all cases the CoV after the application of CCA is lower than in the equivalent data before CCA (light versus dark green bars with respect to each method; Figure 8). Further, the CoV results demonstrate that after application of CCA, the differences between conditions (Uncleaned, PCA-OBS, SSP) are minor, i.e. regardless of the single-trial variation beforehand, the single-trial variation afterwards will be rather similar across conditions (Figure 8). Nevertheless, performing SSP before CCA leads to lower variability as compared to both Uncleaned and PCA-OBS cleaned data, though this effect is only significant when comparing SSP-cleaned to Uncleaned data in the lumbar cord () where cardiac artefacts are most prominent. For exemplary single-trial depictions after the application of cardiac artefact correction methods and CCA as well as grand-average SEP time-courses, see the Supplementary Material.

**Figure 8:**
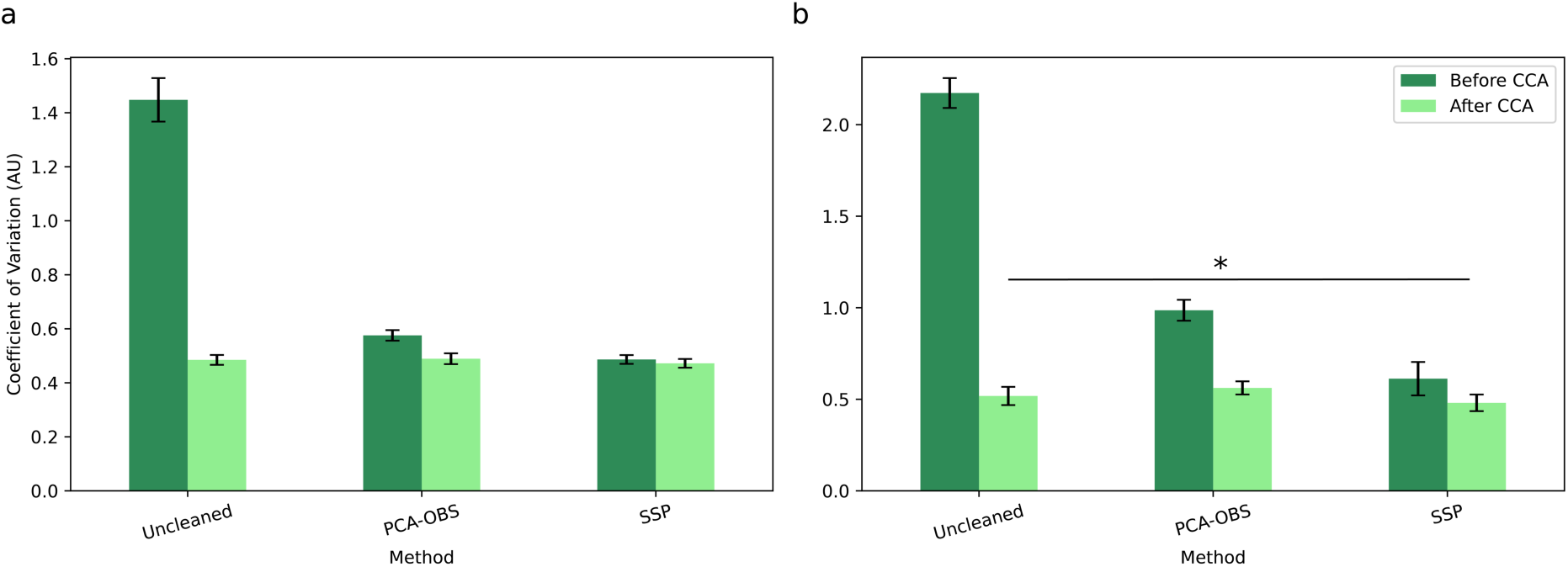
Coefficient of variation results before and after the application of CCA in the cervical spinal cord (a) and the lumbar spinal cord (b). Smaller numbers indicate lower variation across trials. Error bars indicate the standard error of the mean computed across participants. Methods with significant differences (p<0.05) in single-trial variability after the application of CCA are indicated with black lines and asterisks.

**Table 3:** Single-trial coefficient of variation (CoV) and statistical tests for different combinations of methods. The mean of the single trial CoV for all participants, after CCA has been applied (top), with the statistical significance between different methods (bottom), those that reach significance (p<0.05) are shaded green, while those that do not reach significance are shaded yellow.

| Mean single trial variation ± standard error of the mean | | |
| --- | --- | --- |
|  | Median Nerve Stimulation | Tibial Nerve Stimulation |
| Uncleaned + CCA | 0.48 ± 0.02 | 0.52 ± 0.05 |
| PCA-OBS + CCA | 0.49 ± 0.02 | 0.56 ± 0.04 |
| SSP + CCA | 0.47 ± 0.02 | 0.48 ± 0.05 |
| Statistical significance ($p$-values, two-tailed) | | |
| Uncleaned – PCA-OBS | 0.889 | 0.553 |
| Uncleaned – SSP | 0.162 | 0.027 |
| PCA-OBS – SSP | 0.072 | 0.079 |

## 4. Discussion

In this study, we systematically examined the potential of several algorithms to reduce the impact of physiological noise of cardiac origin in spinal cord electrophysiology data. Removing cardiac artefacts is of utmost importance in ESG data, considering that they are between two and three orders of magnitude larger than the evoked potentials of interest. We investigated the impact the application of several common artefact correction methods had both on the cardiac artefact itself and on the somatosensory evoked potentials of interest. These analyses allowed us to determine and compare the strengths and weaknesses of each method in the – until now not investigated – spinal domain and to provide recommendations for their use in specific experimental scenarios.

### 4.1. Overview of each method’s performance

Our investigations suggest that both PCA-OBS and SSP represent effective methods to reduce the influence of cardiac activity on ESG data, with SSP offering a better performance overall, while ICA was seen to be unsuitable for the removal of cardiac activity. Further, we were able to demonstrate that when signal enhancement techniques such as CCA are employed, it is possible to obtain high-fidelity SEPs even in the absence of dedicated cleaning of the cardiac artefact.

#### 4.1.1. PCA-OBS

The application of PCA-OBS more than doubled the SNR of spinal SEPs as compared to Uncleaned data in both the cervical and lumbar spinal cord and was also seen to significantly reduce the impact of the cardiac artefact, e.g. showing a residual artefact intensity of just above one percent. Despite this, the PCA-OBS algorithm can result in sharp voltage deviations at the beginning and end of fitting windows around each heartbeat. While this is not directly of concern when studying evoked responses, it presents an inherent disadvantage when it comes to resting-state recordings (Wang et al., 2021). Thus, we present an updated algorithm which includes the multiplication by a Tukey window (see also Chander et al., 2022) to smooth these so-called edge effects and thereby enable the use of the PCA-OBS algorithm in cases where edge effects present problems for analysis. While this can have a negative effect on the ability to effectively remove the influence of cardiac activity (as evidenced by the worsened residual intensity), it still represents a viable avenue to consider for future research as the INPSR is improved overall.

#### 4.1.2. SSP

The use of SSP to remove the cardiac artefact was most successful for our dataset, with the highest SNR for signals of interest of the methods tested, alongside effective removal of the cardiac artefact as evidenced for example by the residual artefact intensity of 0.5%. Despite a more than four-fold increase in SNR compared to uncleaned data, SSP is seen to reduce the amplitude of the signal as compared to alternatives such as PCA-OBS, though this effect is not uncommon, and has been observed in other studies (Haumann et al., 2016). Additionally, while the number of projectors selected for removal is relatively high (six in our case), this is warranted due to the complex spatial and temporal pattern elicited by cardiac activity; in EEG recordings contaminated by transcranial magnetic stimulation (TMS), six to nine projectors are deemed necessary for successful artefact removal (Mutanen et al., 2016), meaning that it is not unexpected to remove a relatively high number of dimensions for complex artefacts. To avoid unnecessarily removing neural content of interest, we recommend that the number of projectors selected for removal should be optimised for each individual study and subject. Additionally, SSP can offer further advantages over the PCA-OBS approach, as noise not related to cardiac activity located in the artefact subspace can also be effectively attenuated by SSP (Vosskuhl et al., 2020).

#### 4.1.3. ICA

In our investigations, ICA was not capable of reducing the impact of the cardiac artefact while at the same time retaining spinal content of interest. This appears to be related to an apparent over-removal of the cardiac artefact, which has been previously noted in studies attempting to remove the ballistocardiographic artefact in simultaneous EEG-fMRI (Bullock et al., 2021). Despite this, ICA has been successfully applied in invasive spinal electrophysiology recordings to remove cardiac interference (Wang et al., 2021), and in surface electromyography of back muscles (Jiang et al., 2021), though in both cases with different recording arrangements than present here. This indicates that while using automatic component selection via cross-phase trial statistics as performed here results in an over-removal of the cardiac artefact, it is in theory possible to find a critical number of components to effectively remove the cardiac artefact. Further studies may also address the effectiveness of alternative ICA algorithms (e.g. Infomax (Bell and Sejnowski, 1995)) in contrast to the FastICA algorithm employed here. It may also be possible to consider components to remove based on the cardiac activity detected in lateral electrodes along the surface of the back, which should capture the variance of the heartbeat in a more similar capacity to the spinal electrodes of interest, as opposed to the ECG channel itself, though this remains beyond the scope of this work, considering that only very few lateral electrodes are present in the current recording montage.

#### 4.1.4. CCA

As an alternative approach to cardiac artefact removal methods, we presented a method for signal *enhancement* by leveraging a variant of CCA. This method enabled the investigation of SEPs of interest in relation to both median and tibial nerve stimulation, even at the single-trial level, and was shown to be relatively insensitive to pre-cleaning of the cardiac artefact, though a combination with SSP led to further improvement. CCA has previously been shown to be effective for spinal recordings (Nierula et al., 2024), though this is the first time, to the best of our knowledge, that this method has been shown to function well in spinal recordings even in the absence of dedicated cleaning of the cardiac artefact.

### 4.2. Method selection depends on recording setup and research aims

Since PCA-OBS, SSP and CCA all performed well, we now provide recommendations for their selection in specific settings. For removal of the cardiac artefact, in instances where electrode arrays are extensive enough to make use of spatial filtering techniques, we recommend SSP with the number of projectors optimised for each individual study, as also carried out here. However, in cases where small numbers of electrodes (or even single electrodes) are used to record from spinal signals, PCA-OBS offers sufficient performance, and should be the chosen algorithm. As the standard implementation of PCA-OBS can result in the introduction of sharp voltage deviations, it’s use should be limited to the study of evoked potentials and should be avoided in the case of resting-state recordings in the spinal cord. Further improvements may be offered in the cervical cord via the use of anterior re-referencing (Nierula et al., 2024), though this was not implemented here.

Additionally, in cases of task-based recordings with multiple electrodes and no necessity to analyse raw data traces, it is possible to use CCA to obtain high-fidelity somatosensory evoked potentials. This can be pursued without targeted cleaning of the cardiac artefact (as CCA’s performance was sufficient alone), though a combination with SSP is expected to lead to further sensitivity gains. Leveraging spatial filtering approaches like CCA, can enable the extraction of more robust spinal SEPs and will be particularly useful in instances where large trial counts are unsuitable, for example, in pain research. Here, it was also possible to obtain clear somatosensory evoked potentials at the single trial level, which opens further paths for analysis, including the investigation of the fluctuation of response amplitudes across different processing levels of the peripheral and central nervous system (Nierula et al., 2024).

### 4.3. Advantages of cardiac artefact denoising over classical approaches

Beyond enabling single trial investigations, the cardiac denoising methods included in this investigation have significant advantages over classical approaches to limit the impact of the cardiac artefact. As mentioned previously, these approaches include, i) cardiac gated stimulation, ii) trial averaging, and iii) high-pass filtering. Cardiac-gated stimulation involves stimulating only during artefact-free periods of the cardiac cycle (Cracco, 1973), and while it is an effective means to eliminate the cardiac artefact, it does not allow for studies where cardiac-somatosensory interactions are of interest and stimulation must be distributed across the cardiac cycle (Al et al., 2020). In contrast, the noise reduction methods employed in this investigation do not preclude examining such interactions. Further, for successful trial averaging, typically ~2000 trials are necessary (Cruccu et al., 2008). However, modern cognitive neuroscience paradigms often consist of conditions with low trial numbers (e.g. deviance detection designs in the context of predictive processing; (Tabas et al., 2020)), therefore, the use of cardiac denoising methods that provide high-fidelity signals even with low trial counts, such as exhibited here, are vital. Finally, ESG studies often filter out content below 30Hz (Lastimosa et al., 1982) which helps to remove frequency content associated with cardiac activity. While high-pass filtering is not problematic in the investigation of spinal somatosensory evoked potentials, it is undesirable in the case of resting-state recordings, where frequencies of interest can lie below 30Hz (Wang et al., 2021). The methods shown here are proven effective even without such extensive high-pass filtering, and are thus more suitable for studies involving resting state recordings.

### 4.4. Alternative approaches

While this study did not exhaustively examine all possible methods to reduce the impact of the cardiac artefact in spinal electrophysiology recordings, to the best of our knowledge it represents the first study to evaluate how well common methods for artefact removal perform in the spinal domain.

Alternative approaches not considered in the present study – but which have had varying success in cardiac artefact removal in alternative domains – include for example template-based subtraction (Chander et al., 2022), empirical mode decomposition in combination with principal component analysis (Javed et al., 2017), deep learning (McIntosh et al., 2021), and harmonic regression approaches (Krishnaswamy et al., 2016). Additionally, given the promising results of ICA in terms of cardiac artefact removal, and the ability of SSP to improve the SNR of our signal of interest, it may be possible in the future to create a hybrid method that balances the advantages of each approach and can more fully remove the impact of the cardiac artefact without compromising the quality of the signal of interest (this has been attempted with some success with respect to blink correction in EEG recordings (Korpela et al., 2012)).

While signal space projection (SSP) was chosen in this study due to its ability to remove cardiac artefacts from EEG and MEG data, a related method termed dual signal subspace projection (DSSP) (Sekihara et al., 2016) has recently been introduced in the magnetospinography community for the removal of stimulation artefacts (Akaza et al., 2021). Despite promising initial results, to the best of our knowledge there have neither been attempts to extend this algorithm towards the removal of cardiac activity nor has it been applied in the context of electroencephalographic or electrospinographic data and thus remains a candidate algorithm for future investigations.

Although CCA was examined in this study in the context of enhancing the somatosensory evoked potentials of interest, it is also possible to instead use CCA to capture the pattern of the cardiac artefact and focus directly on its removal (Assecondi et al., 2010, 2009). Also, each of the top four components identified by CCA were manually examined, and only the one best matching the expected time course, spatial topography and having the best single trial representation was selected, as previously performed (Stephani et al., 2020). However, other studies have suggested that a combination of multiple components can provide a better result than the top component alone (de Cheveigné et al., 2018), and this is a potential avenue for further study, alongside an automated selection of the best component to eliminate researcher degrees of freedom (Simmons et al., 2011).

Ultimately, the difficulty in removing the cardiac artefact from spinal electrophysiology recordings lies in its large amplitude relative to the spinal activity of interest, and the spatial and temporal non-stationary characteristics of cardiac activity (i.e. there can be changes in the timing and shape of heartbeat occurrences over time). As all methods tested here assume temporal stationarity, to achieve near complete removal of the cardiac artefact, it may become necessary to look beyond current methods and develop improved approaches tailored to the specific and dynamic activity of the heart.

### 4.5. Limitations

There are several limitations in the present study worthy of mention. First, it is important to note that there was no significant locking of the cardiac cycle to the somatosensory stimulation, and in cases of such temporal locking of these events, the cardiac artefact removal methods examined here may perform sub-optimally. Second, while this study presents an analysis of several common algorithms used to remove cardiac interference in cortical recordings, it is not an exhaustive study of all possible cardiac artefact removal methods and thus there might exist even better methods to remove the cardiac artefact in spinal electrophysiological recordings. For example, improvements could be made by considering the impact of respiration on the cardiac cycle. Finally, since the results of this study are specific to our high-density recording montage and may vary in alternative recording set-ups, it is important to consider the specifics of each study in order to select the appropriate algorithm for each individual use case.

### 4.6. Conclusion

Experimental research in non-invasive electrophysiology of the human spinal cord is currently only being performed by few research groups and there has thus far been no focus on determining effective methods to reduce the effect of the cardiac artefact that limits the utility of spinal surface electrodes. This study evaluated some of the most common artefact reduction techniques in imaging neuroscience and examined their efficacy at removing the cardiac artefact in electrospinography data. When electrode arrays are extensive enough to allow for spatial filtering, SSP is recommended, while in cases where the recording montage does not permit this, PCA-OBS and its variations should be considered depending on the aims of study. Further, using CCA allows for obtaining high-fidelity somatosensory evoked potentials, even in the absence of dedicated cleaning methods. Taken together, our investigations provide viable approaches for increasing the robustness and sensitivity of electrospinography across different recording set-ups as well as different experimental scenarios, such as task-based or resting-state investigations.

## Data and code availability

The underlying data and code are openly available via OpenNeuro (https://openneuro.org/datasets/ds004388) and GitHub (https://github.com/eippertlab/cardiac-artefact-removal), respectively.

## Ethics

All participants gave written informed consent. The study was approved by the Ethics Committee of the Medical Faculty of the University of Leipzig.

## Author contributions

Author contributions are listed alphabetically according to CRediT taxonomy (https://credit.niso.org).

Conceptualization: EB, FE, BM, BN, VN, TS

Data curation: BN

Formal analysis: EB

Funding acquisition: FE

Investigation: EB, BN

Methodology: EB, FE, BM

Project administration: EB, FE

Resources: FE, VN

Software: EB, BN

Supervision: FE, BM, VN

Visualization: EB

Writing – original draft: EB, FE

Writing – review & editing: EB, FE, BM, BN, TS, VN

## Acknowledgements

We would like to thank all volunteers who participated in this study. Additionally, we want to thank everyone who assisted in data acquisition.

## References

Abraira, V.E., Kuehn, E.D., Chirila, A.M., Springel, M.W., Toliver, A.A., Zimmerman, A.L., Orefice, L.L., Boyle, K.A., Bai, L., Song, B.J., Bashista, K.A., O’Neill, T.G., Zhuo, J., Tsan, C., Hoynoski, J., Rutlin, M., Kus, L., Niederkofler, V., Watanabe, M., Dymecki, S.M., Nelson, S.B., Heintz, N., Hughes, D.I., Ginty, D.D., 2017. The Cellular and Synaptic Architecture of the Mechanosensory Dorsal Horn. Cell 168, 295–310.e19. 10.1016/j.cell.2016.12.010

Adachi, Y., Kawabata, S., Hashimoto, J., Okada, Y., Naijo, Y., Watanabe, T., Miyano, Y., Uehara, G., 2021. Multichannel SQUID Magnetoneurograph System for Functional Imaging of Spinal Cords and Peripheral Nerves. IEEE Trans. Appl. Supercond. 31, 1–5. 10.1109/TASC.2021.3056492

Ahuja, C.S., Wilson, J.R., Nori, S., Kotter, M.R.N., Druschel, C., Curt, A., Fehlings, M.G., 2017. Traumatic spinal cord injury. Nat. Rev. Dis. Primer 3, 17018. 10.1038/nrdp.2017.18

Akaza, M., Kawabata, S., Ozaki, I., Miyano, Y., Watanabe, T., Adachi, Y., Sekihara, K., Sumi, Y., Yokota, T., 2021. Noninvasive measurement of sensory action currents in the cervical cord by magnetospinography. Clin. Neurophysiol. 132, 382–391. 10.1016/j.clinph.2020.11.029

Al, E., Iliopoulos, F., Forschack, N., Nierhaus, T., Grund, M., Motyka, P., Gaebler, M., Nikulin, V.V., Villringer, A., 2020. Heart–brain interactions shape somatosensory perception and evoked potentials. Proc. Natl. Acad. Sci. 117, 10575–10584. 10.1073/pnas.1915629117

Assecondi, S., Hallez, H., Staelens, S., Bianchi, A.M., Huiskamp, G.M., Lemahieu, I., 2009. Removal of the ballistocardiographic artifact from EEG–fMRI data: a canonical correlation approach. Phys. Med. Biol. 54, 1673–1689. 10.1088/0031-9155/54/6/018

Assecondi, S., Vanderperren, K., Novitskiy, N., Ramautar, J.R., Fias, W., Staelens, S., Stiers, P., Sunaert, S., Van Huffel, S., Lemahieu, I., 2010. Effect of the static magnetic field of the MR-scanner on ERPs: Evaluation of visual, cognitive and motor potentials. Clin. Neurophysiol. 121, 672–685. 10.1016/j.clinph.2009.12.032

Bell, A.J., Sejnowski, T.J., 1995. An Information-Maximization Approach to Blind Separation and Blind Deconvolution. Neural Comput. 7, 1129–1159. 10.1162/neco.1995.7.6.1129

Bullock, M., Jackson, G.D., Abbott, D.F., 2021. Artifact Reduction in Simultaneous EEG-fMRI: A Systematic Review of Methods and Contemporary Usage. Front. Neurol. 12, 622719. 10.3389/fneur.2021.622719

Chander, B.S., Deliano, M., Azañón, E., Büntjen, L., Stenner, M.-P., 2022. Non-invasive recording of high-frequency signals from the human spinal cord. NeuroImage 253, 119050. 10.1016/j.neuroimage.2022.119050

Ciccarelli, O., Cohen, J.A., Reingold, S.C., Weinshenker, B.G., Amato, M.P., Banwell, B., Barkhof, F., Bebo, B., Becher, B., Bethoux, F., Brandt, A., Brownlee, W., Calabresi, P., Chatway, J., Chien, C., Chitnis, T., Ciccarelli, O., Cohen, J., Comi, G., Correale, J., De Sèze, J., De Stefano, N., Fazekas, F., Flanagan, E., Freedman, M., Fujihara, K., Galetta, S., Goldman, M., Greenberg, B., Hartung, H.-P., Hemmer, B., Henning, A., Izbudak, I., Kappos, L., Lassmann, H., Laule, C., Levy, M., Lublin, F., Lucchinetti, C., Lukas, C., Marrie, R.A., Miller, A., Miller, D., Montalban, X., Mowry, E., Ourselin, S., Paul, F., Pelletier, D., Ranjeva, J.-P., Reich, D., Reingold, S., Rocca, M.A., Rovira, A., Schlaerger, R., Soelberg Sorensen, P., Sormani, M., Stuve, O., Thompson, A., Tintoré, M., Traboulsee, A., Trapp, B., Trojano, M., Uitdehaag, B., Vukusic, S., Waubant, E., Weinshenker, B., Wheeler-Kingshott, C.G., Xu, J., 2019. Spinal cord involvement in multiple sclerosis and neuromyelitis optica spectrum disorders. Lancet Neurol. 18, 185–197. 10.1016/S1474-4422(18)30460-5

Cracco, R.Q., 1973. Spinal evoked response: Peripheral nerve stimulation in man. Electroencephalogr. Clin. Neurophysiol. 35, 379–386. 10.1016/0013-4694(73)90195-8

Cracco, R.Q., Cracco, J.B., Anziska, B.J., 1979. Somatosensory Evoked Potentials in Man: Cerebral, Subcortical, Spinal, and Peripheral Nerve Potentials. Am. J. EEG Technol. 19, 59–81. 10.1080/00029238.1979.11079967

Cruccu, G., Aminoff, M.J., Curio, G., Guerit, J.M., Kakigi, R., Mauguiere, F., Rossini, P.M., Treede, R.-D., Garcia-Larrea, L., 2008. Recommendations for the clinical use of somatosensory-evoked potentials. Clin. Neurophysiol. 119, 1705–1719. 10.1016/j.clinph.2008.03.016

Dammers, J., Schiek, M., Boers, F., Silex, C., Zvyagintsev, M., Pietrzyk, U., Mathiak, K., 2008. Integration of Amplitude and Phase Statistics for Complete Artifact Removal in Independent Components of Neuromagnetic Recordings. IEEE Trans. Biomed. Eng. 55, 2353–2362. 10.1109/TBME.2008.926677

de Cheveigné, A., Wong, D.D.E., Di Liberto, G.M., Hjortkjær, J., Slaney, M., Lalor, E., 2018. Decoding the auditory brain with canonical component analysis. NeuroImage 172, 206–216. 10.1016/j.neuroimage.2018.01.033

Desmedt, J.E., Cheron, G., 1982. Somatosensory Evoked Potentials in Man: Subcortical and Cortical Components and their Neural Basis. Ann. N. Y. Acad. Sci. 388, 388–410. 10.1111/j.1749-6632.1982.tb50804.x

Fedele, T., Scheer, H.-J., Burghoff, M., Waterstraat, G., Nikulin, V.V., Curio, G., 2013. Distinction between added-energy and phase-resetting mechanisms in non-invasively detected somatosensory evoked responses, in: 2013 35th Annual International Conference of the IEEE Engineering in Medicine and Biology Society (EMBC). Presented at the 2013 35th Annual International Conference of the IEEE Engineering in Medicine and Biology Society (EMBC), IEEE, Osaka, pp. 1688–1691. 10.1109/EMBC.2013.6609843

Gijsen, S., Grundei, M., Lange, R.T., Ostwald, D., Blankenburg, F., 2021. Neural surprise in somatosensory Bayesian learning. PLOS Comput. Biol. 17, e1008068. 10.1371/journal.pcbi.1008068

Gramfort, A., 2013. MEG and EEG data analysis with MNE-Python. Front. Neurosci. 7. 10.3389/fnins.2013.00267

Haufe, S., Meinecke, F., Görgen, K., Dähne, S., Haynes, J.-D., Blankertz, B., Bießmann, F., 2014. On the interpretation of weight vectors of linear models in multivariate neuroimaging. NeuroImage 87, 96–110. 10.1016/j.neuroimage.2013.10.067

Haumann, N.T., Parkkonen, L., Kliuchko, M., Vuust, P., Brattico, E., 2016. Comparing the Performance of Popular MEG/EEG Artifact Correction Methods in an Evoked-Response Study. Comput. Intell. Neurosci. 2016, 1–10. 10.1155/2016/7489108

Hochman, S., 2007. Spinal cord. Curr. Biol. 17, R950–R955. 10.1016/j.cub.2007.10.014

Hyvarinen, A., 1999. Fast and robust fixed-point algorithms for independent component analysis. IEEE Trans. Neural Netw. 10, 626–634. 10.1109/72.761722

Javed, E., Faye, I., Malik, A.S., Abdullah, J.M., 2017. Removal of BCG artefact from concurrent fMRI-EEG recordings based on EMD and PCA. J. Neurosci. Methods 291, 150–165. 10.1016/j.jneumeth.2017.08.020

Jiang, N., Wang, L., Huang, Z., Li, G., 2021. Mapping Responses of Lumbar Paravertebral Muscles to Single-Pulse Cortical TMS Using High-Density Surface Electromyography. IEEE Trans. Neural Syst. Rehabil. Eng. 29, 831–840. 10.1109/TNSRE.2021.3076095

Jones, S.J., Small, D.G., 1978. Spinal and sub-cortical evoked potentials following stimulation of the posterior tibial nerve in man. Electroencephalogr. Clin. Neurophysiol. 44, 299–306. 10.1016/0013-4694(78)90305-X

Kinany, N., Pirondini, E., Micera, S., Van De Ville, D., 2022. Spinal Cord fMRI: A New Window into the Central Nervous System. The Neuroscientist 107385842211018. 10.1177/10738584221101827

Korpela, J., Vigario, R., Huotilainen, M., 2012. The effect of automatic blink correction on auditory evoked potentials, in: 2012 Annual International Conference of the IEEE Engineering in Medicine and Biology Society. Presented at the 2012 34th Annual International Conference of the IEEE Engineering in Medicine and Biology Society (EMBC), IEEE, San Diego, CA, pp. 625–628. 10.1109/EMBC.2012.6346009

Krishnaswamy, P., Bonmassar, G., Poulsen, C., Pierce, E.T., Purdon, P.L., Brown, E.N., 2016. Reference-free removal of EEG-fMRI ballistocardiogram artifacts with harmonic regression. NeuroImage 128, 398–412. 10.1016/j.neuroimage.2015.06.088

Kuner, R., Flor, H., 2017. Structural plasticity and reorganisation in chronic pain. Nat. Rev. Neurosci. 18, 20–30. 10.1038/nrn.2016.162

Landelle, C., Lungu, O., Vahdat, S., Kavounoudias, A., Marchand-Pauvert, V., De Leener, B., Doyon, J., 2021. Investigating the human spinal sensorimotor pathways through functional magnetic resonance imaging. NeuroImage 245, 118684. 10.1016/j.neuroimage.2021.118684

Lastimosa, A.C.B., Bass, N.H., Stanback, K., Norvell, E.E., 1982. Lumbar spinal cord and early cortical evoked potentials after tibial nerve stimulation: Effects of stature on normative data. Electroencephalogr. Clin. Neurophysiol. 54, 499–507. 10.1016/0013-4694(82)90035-9

Mantini, D., Perrucci, M.G., Del Gratta, C., Romani, G.L., Corbetta, M., 2007. Electrophysiological signatures of resting state networks in the human brain. Proc. Natl. Acad. Sci. 104, 13170– 13175. 10.1073/pnas.0700668104

Mardell, L.C., O’Neill, G.C., Tierney, T.M., Timms, R.C., Zich, C., Barnes, G.R., Bestmann, S., 2022. Concurrent spinal and brain imaging with optically pumped magnetometers (preprint). Neuroscience. 10.1101/2022.05.12.491623

Marino, M., Liu, Q., Koudelka, V., Porcaro, C., Hlinka, J., Wenderoth, N., Mantini, D., 2018. Adaptive optimal basis set for BCG artifact removal in simultaneous EEG-fMRI. Sci. Rep. 8, 8902. 10.1038/s41598-018-27187-6

Mauguière, F., 2000. Anatomic Origin of the Cervical N13 Potential Evoked by Upper Extremity Stimulation: J. Clin. Neurophysiol. 17, 236–245. 10.1097/00004691-200005000-00002

McIntosh, J.R., Yao, J., Hong, L., Faller, J., Sajda, P., 2021. Ballistocardiogram Artifact Reduction in Simultaneous EEG-fMRI Using Deep Learning. IEEE Trans. Biomed. Eng. 68, 78–89. 10.1109/TBME.2020.3004548

Mutanen, T.P., Kukkonen, M., Nieminen, J.O., Stenroos, M., Sarvas, J., Ilmoniemi, R.J., 2016. Recovering TMS-evoked EEG responses masked by muscle artifacts. NeuroImage 139, 157–166. 10.1016/j.neuroimage.2016.05.028

Niazy, R.K., Beckmann, C.F., Iannetti, G.D., Brady, J.M., Smith, S.M., 2005. Removal of FMRI environment artifacts from EEG data using optimal basis sets. NeuroImage 28, 720–737. 10.1016/j.neuroimage.2005.06.067

Nichols, T.E., Holmes, A.P., 2002. Nonparametric permutation tests for functional neuroimaging: A primer with examples. Hum. Brain Mapp. 15, 1–25. 10.1002/hbm.1058

Nierula, B., Stephani, T., Bailey, E., Kaptan, M., Pohle, L.-M., Horn, U., Mouraux, A., Maess, B., Villringer, A., Curio, G., Nikulin, V.V., Eippert, F., 2024. A multi-channel electrophysiology approach to non-invasively and precisely record human spinal cord activity. 10.1101/2022.12.05.519148

Paixão, S., Loschek, L., Gaitanos, L., Alcalà Morales, P., Goulding, M., Klein, R., 2019. Identification of Spinal Neurons Contributing to the Dorsal Column Projection Mediating Fine Touch and Corrective Motor Movements. Neuron 104, 749–764.e6. 10.1016/j.neuron.2019.08.029

Sekihara, K., Kawabata, Y., Ushio, S., Sumiya, S., Kawabata, S., Adachi, Y., Nagarajan, S.S., 2016. Dual signal subspace projection (DSSP): a novel algorithm for removing large interference in biomagnetic measurements. J. Neural Eng. 13, 036007. 10.1088/1741-2560/13/3/036007

Simmons, J.P., Nelson, L.D., Simonsohn, U., 2011. False-Positive Psychology: Undisclosed Flexibility in Data Collection and Analysis Allows Presenting Anything as Significant. Psychol. Sci. 22, 1359– 1366. 10.1177/0956797611417632

Stephani, T., Waterstraat, G., Haufe, S., Curio, G., Villringer, A., Nikulin, V.V., 2020. Temporal Signatures of Criticality in Human Cortical Excitability as Probed by Early Somatosensory Responses. J. Neurosci. 40, 6572–6583. 10.1523/JNEUROSCI.0241-20.2020

Sumiya, S., Kawabata, S., Hoshino, Y., Adachi, Y., Sekihara, K., Tomizawa, S., Tomori, M., Ishii, S., Sakaki, K., Ukegawa, D., Ushio, S., Watanabe, T., Okawa, A., 2017. Magnetospinography visualizes electrophysiological activity in the cervical spinal cord. Sci. Rep. 7, 2192. 10.1038/s41598-017-02406-8

Tabas, A., Mihai, G., Kiebel, S., Trampel, R., von Kriegstein, K., 2020. Abstract rules drive adaptation in the subcortical sensory pathway. eLife 9, e64501. 10.7554/eLife.64501

Uusitalo, M.A., Ilmoniemi, R.J., 1997. Signal-space projection method for separating MEG or EEG into components. Med. Biol. Eng. Comput. 35, 135–140. 10.1007/BF02534144

Vosskuhl, J., Mutanen, T.P., Neuling, T., Ilmoniemi, R.J., Herrmann, C.S., 2020. Signal-Space Projection Suppresses the tACS Artifact in EEG Recordings. Front. Hum. Neurosci. 14, 536070. 10.3389/fnhum.2020.536070

Wang, F., Zhang, L., Yue, L., Zeng, Y., Zhao, Q., Gong, Q., Zhang, J., Liu, D., Luo, X., Xia, X., Wan, L., Hu, L., 2021. A novel method to simultaneously record spinal cord electrophysiology and electroencephalography signals. NeuroImage 232, 117892. 10.1016/j.neuroimage.2021.117892

Waterstraat, G., Fedele, T., Burghoff, M., Scheer, H.-J., Curio, G., 2015. Recording human cortical population spikes non-invasively – An EEG tutorial. J. Neurosci. Methods 250, 74–84. 10.1016/j.jneumeth.2014.08.013

Yamada, T., Kimura, J., Nitz, D.M., 1980. Short latency somatosensory evoked potentials following median nerve stimulation in man. Electroencephalogr. Clin. Neurophysiol. 48, 367–376. 10.1016/0013-4694(80)90129-7

