## Supplementary Material for "Evaluating noise correction approaches for non-invasive electrophysiology of the human spinal cord"

### 1. Choosing the number of projectors for signal space projection

In order to choose the number of projectors to remove the cardiac artefact in the case of signal space projection (SSP), each number of projectors from 1 to 20 was tested. Following this, the signal-to-noise ratio (SNR) of the somatosensory evoked potential (SEP) of interest, alongside the residual intensity (RI) and improved normalised power spectrum ratio (INPSR) of the cardiac artefact was computed across all participants. This enabled a direct comparison between the number of projectors chosen and the subsequent effect on both the SEPs and the cardiac artefact to make an informed decision. The results in terms of median nerve stimulation in the cervical spinal cord can be seen in Table S1, while the equivalent results for tibial nerve stimulation in the lumbar spinal cord can be seen in Table S2. Based on these results, six projectors were chosen in both cases, as this presented the optimal balance between removing the cardiac artefact and retaining a high SNR of the SEPs: the SNR suffers minimally when moving from the number of projectors with the strongest SNR (five in the cervical cord, four in the lumbar cord) to six projectors, while the RI and INPSR substantially improve. Beyond six projectors, the SNR begins to decline rapidly, while the benefits in terms of the RI and INPSR slowly plateau.

*Table S1: Residual intensity (RI), improved normalised power spectrum ratio (INPSR) of the cardiac artefact, and the signal-to-noise ratio (SNR) of the SEPs for median nerve stimulation in the cervical spinal cord for differing numbers of projectors, calculated across all participants. The highlighted row depicts the results for the chosen number of projectors.*

| NUMBER OF PROJECTORS | RI | INPSR | SNR |
|---|---|---|---|
| **1** | 48.12% | 7.12E+00 | 2.75 |
| **2** | 9.25% | 2.96E+02 | 8.58 |
| **3** | 3.22% | 1.12E+03 | 11.69 |
| **4** | 0.85% | 2.64E+03 | 14.20 |
| **5** | 0.57% | 3.05E+03 | 14.37 |
| **6** | 0.43% | 3.81E+03 | 14.30 |
| **7** | 0.37% | 4.30E+03 | 14.05 |
| **8** | 0.28% | 5.38E+03 | 13.90 |
| **9** | 0.23% | 5.57E+03 | 12.76 |
| **10** | 0.20% | 6.53E+03 | 12.51 |
| **11** | 0.17% | 8.04E+03 | 12.00 |
| **12** | 0.15% | 9.13E+03 | 10.47 |
| **13** | 0.13% | 8.37E+03 | 9.68 |
| **14** | 0.11% | 9.09E+03 | 9.74 |
| **15** | 0.11% | 9.74E+03 | 8.99 |
| **16** | 0.10% | 1.13E+04 | 8.55 |
| **17** | 0.10% | 1.13E+04 | 8.14 |
| **18** | 0.09% | 1.23E+04 | 7.76 |
| **19** | 0.09% | 1.26E+04 | 7.13 |
| **20** | 0.08% | 1.34E+04 | 7.30 |

*Table S2: Residual intensity (RI), improved normalised power spectrum ratio (INPSR) of the cardiac artefact, and the signal-to-noise ratio (SNR) of the SEPs for tibial nerve stimulation in the lumbar spinal cord for differing numbers of projectors, calculated across all participants. The highlighted row depicts the results for the chosen number of projectors.*

| NUMBER OF PROJECTORS | RI | INPSR | SNR |
|---|---|---|---|
| **1** | 8.86% | 5.01E+02 | 4.22 |
| **2** | 3.50% | 1.80E+03 | 5.59 |
| **3** | 2.12% | 5.01E+03 | 7.41 |
| **4** | 0.99% | 1.22E+04 | 10.46 |
| **5** | 0.48% | 2.13E+04 | 9.92 |
| **6** | 0.29% | 2.90E+04 | 9.80 |
| **7** | 0.20% | 3.31E+04 | 8.00 |
| **8** | 0.14% | 4.13E+04 | 7.47 |
| **9** | 0.11% | 4.32E+04 | 6.98 |
| **10** | 0.10% | 4.01E+04 | 6.07 |
| **11** | 0.09% | 3.51E+04 | 5.40 |
| **12** | 0.08% | 3.81E+04 | 4.88 |
| **13** | 0.07% | 4.14E+04 | 4.15 |
| **14** | 0.07% | 3.83E+04 | 4.01 |
| **15** | 0.06% | 4.18E+04 | 4.10 |
| **16** | 0.06% | 4.63E+04 | 4.15 |
| **17** | 0.05% | 5.23E+04 | 4.21 |
| **18** | 0.05% | 5.86E+04 | 4.09 |
| **19** | 0.05% | 6.08E+04 | 4.18 |
| **20** | 0.04% | 6.68E+04 | 3.76 |

#### 2. Single participant time-courses

The single participant plots in this section supplement the group-level results, with Figure S1 depicting the effect on the cardiac artefact itself and Figure S2 showing the SEP time courses in relation to alternative cleaning pipelines, for a single participant (sub-010).

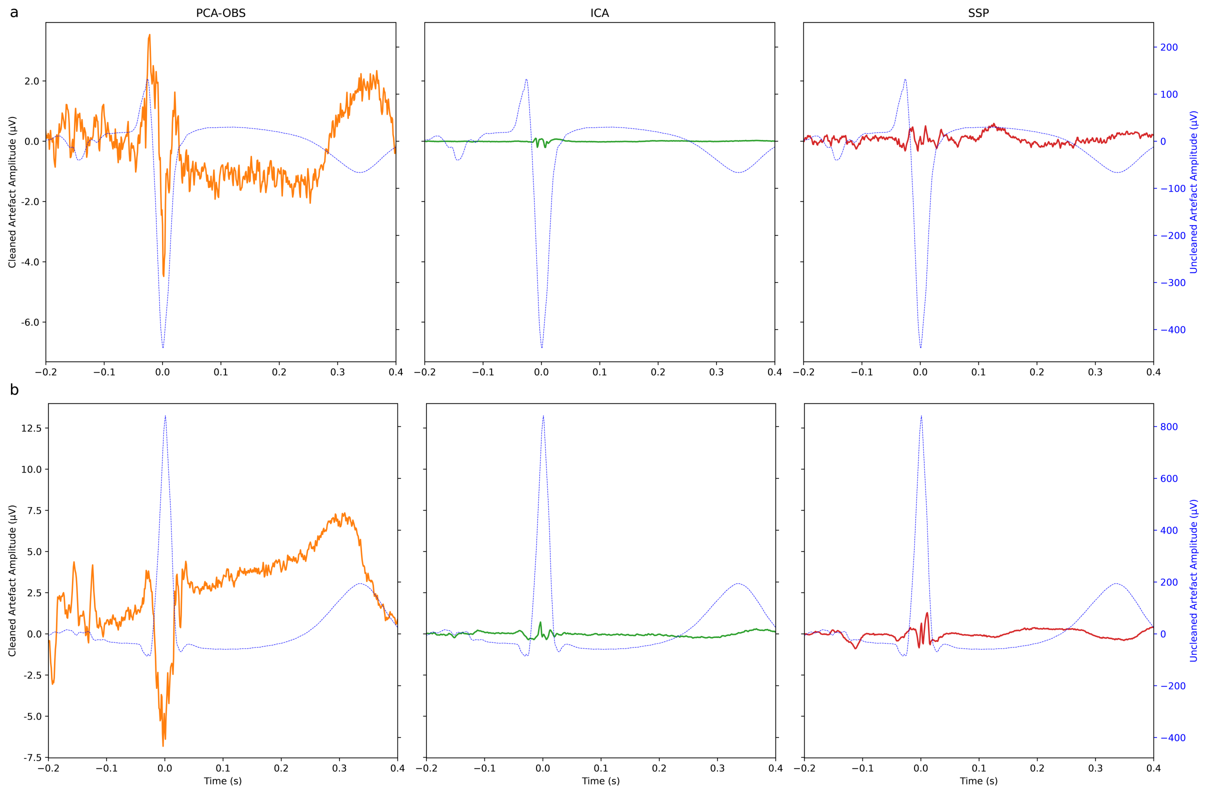


*Figure S1: Single-participant (sub-010) artefact time-courses in the cervical spinal cord (a) and the lumbar spinal cord (b) after application of different cleaning methods, with artefact time-course for Uncleaned data in the background (blue trace).*

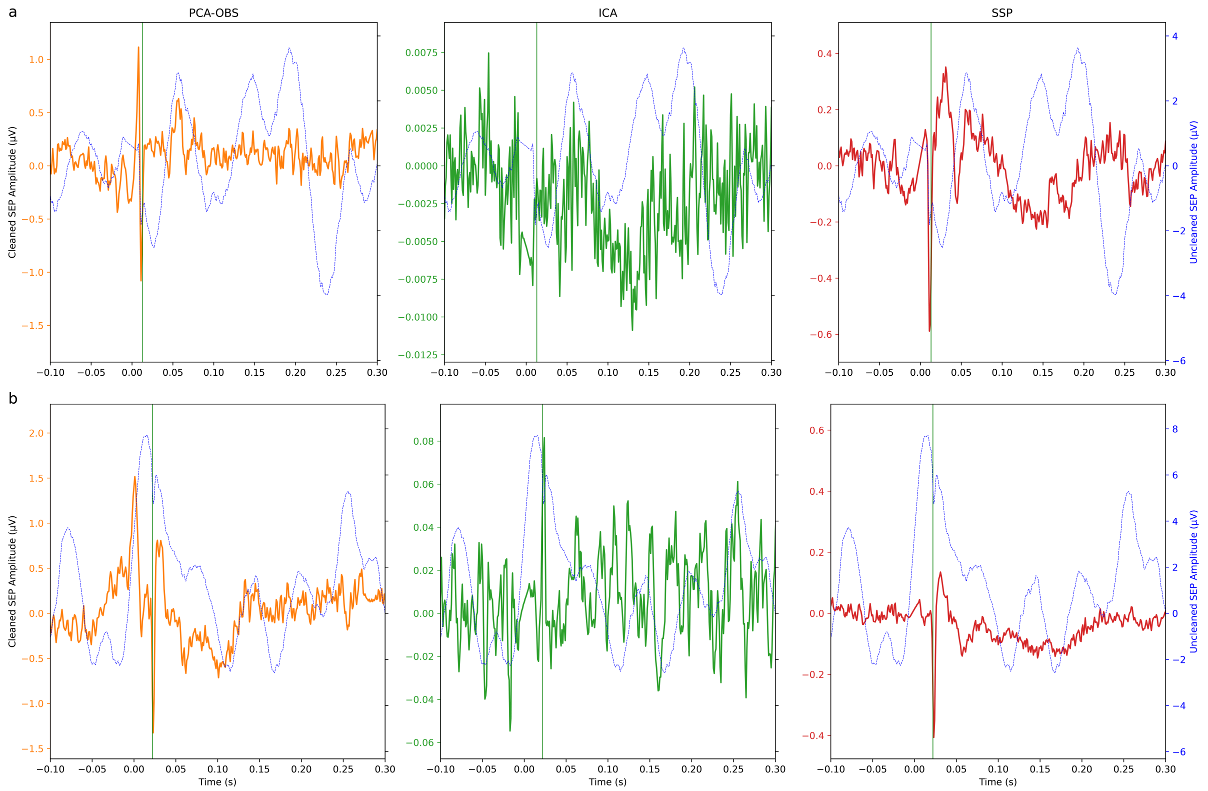


*Figure S2: Single-participant (sub-010) SEP time course in the cervical spinal cord (a) and the lumbar spinal cord (b) after application of different cleaning methods, with SEP time-course for Uncleaned data in the background (blue trace).*

### 3. The effect of a Tukey window with the PCA-OBS algorithm

As previously noted, the PCA-OBS algorithm can introduce sharp voltage deviations at the beginning or end of artefact fitting windows, which is particularly damaging in the case of resting state recordings. To address this, the ability of a Tukey window to reduce this effect was examined visually in a trifold manner: by i) probing the ability to reduce edge effects apparent in the raw data traces, ii) studying the effect on the group-level cardiac artefact, and iii) considering the impact on the group-level somatosensory evoked potentials of interest.

Figure S3 demonstrates the effect of a Tukey window on the raw data traces of a single participant (sub-020), where the original PCA-OBS algorithm is seen to leave sharp deviations in voltage at the edges of fitting windows of the artefact. The updated algorithm that includes the multiplication by a Tukey window is able to effectively smooth these rifts, though it does introduce low-frequency valleys in their place.

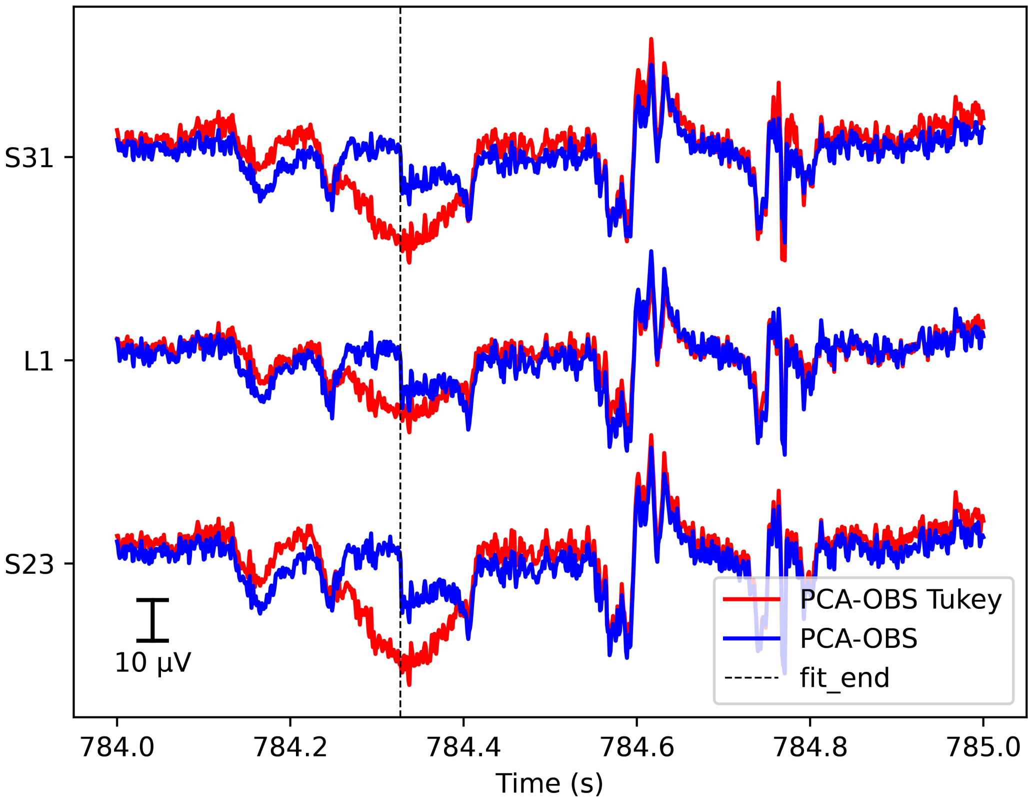


*Figure S3: The effect of the Tukey window (red) on raw data traces from 3 lumbar spinal channels demonstrating the effect in a single participant (sub-020) alongside the original PCA-OBS algorithm (blue), fit_end marks the end of a fitting window about a heartbeat occurrence.*

The effect of the Tukey window modification as compared to the original PCA-OBS algorithm was then examined at the group-level, with Figure S4 exhibiting the difference in terms of the grand averaged cardiac artefact after cleaning (top), and the grand average SEPs of interest (bottom). Evident from the grand average cardiac artefact is the effect of the Tukey window pulling the edges of each fitted artefact to 0V meaning that when the fitted artefact is subtracted from each heartbeat occurrence it removes very little of the original artefact towards the edges of the fitting window (>0.3s). On the other hand, there is a very minimal impact of the Tukey window on the somatosensory evoked potentials; the latency of the evoked potentials of interest remains unchanged between the PCA-OBS and PCA-OBS Tukey cleaned data, with very minor differences in the amplitude seen visually.

These results are supported by the metrics computed, as seen in Table S3. The resulting metrics related to the effect of cleaning on the cardiac artefact reveal a mixed performance regarding the application of a Tukey window – the INPSR is slightly improved, while the RI is negatively affected by the application of a Tukey window. The effect on the somatosensory evoked potentials of interest, as seen via the SNR, is almost negligible. Given the mixed results to the application of the Tukey window, choosing whether or not to implement this additional step will depend on the specific aims of the research and how important it is to avoid the sharp voltage deviations introduced by PCA-OBS.

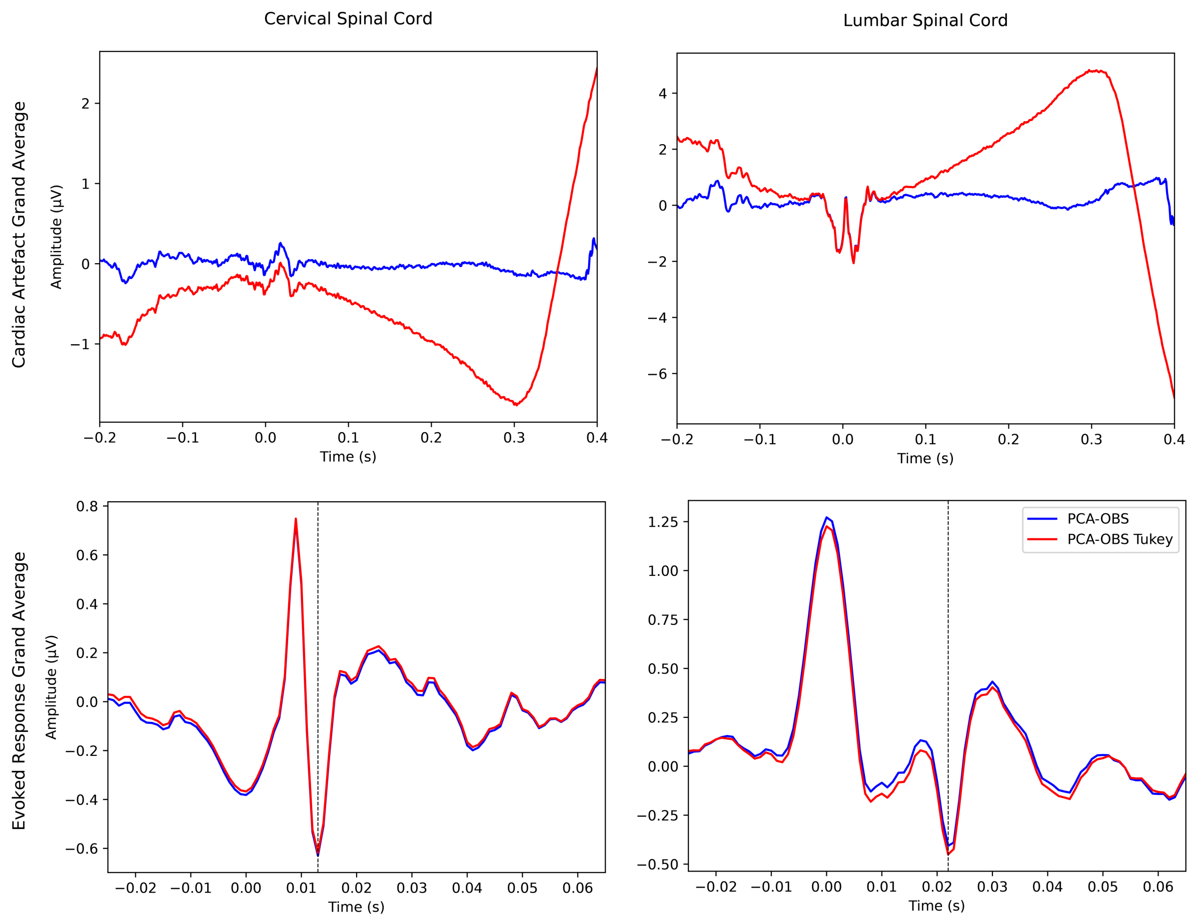


*Figure S4: The top row reveals the effect of the original PCA-OBS algorithm and the modified PCA-OBS Tukey algorithm on the group-level cardiac artefact after cleaning in the cervical spinal cord (left) and the lumbar spinal cord (right). At the R-peak (0s), there is a minimal impact of the Tukey window, but at the edges (beyond 0.3s) the effect of the Tukey window pulling the fitted artefact towards 0V and thus failing to effectively remove this region of the cardiac artefact is apparent. The bottom row then demonstrates the difference between the original PCA-OBS algorithm and the PCA-OBS algorithm modified to include a Tukey window with respect to their effect on the somatosensory evoked potentials of interest, with the expected latency marked by a red line for median nerve stimulation (left) at 13ms, and tibial nerve stimulation (right) at 22ms.*

*Table S3: The results in terms of residual intensity (RI), improved normalised power spectrum ratio (INPSR) of the cardiac artefact, and the signal-to-noise ratio (SNR) of the SEPs for both median and tibial nerve stimulation, calculated across all participants, for the original PCA-OBS algorithm, and the PCA-OBS algorithm including a Tukey window.*

| | **RI** | | **INPSR** | | **SNR** | |
|---|---|---|---|---|---|---|
| | Median | Tibial | Median | Tibial | Median | Tibial |
| **PCA-OBS** | 0.6884% | 1.1620% | 455.55 | 37.40 | 7.6733 | 4.6456 |
| **PCA-OBS TUKEY** | 2.9522% | 4.3519% | 647.55 | 41.84 | 7.6330 | 4.9273 |

#### 4. The effect of PCA-OBS and SSP on low-frequency oscillations

The SEP time-courses after cleaning with PCA-OBS and SSP show severely reduced low-frequency content. To demonstrate the reason this occurs, Figure S5 shows the time courses and power spectra, for both the cervical and lumbar spinal cord in a single participant (sub-020), before and after cleaning of the cardiac artefact via PCA-OBS and SSP. These show that the striking decrease in low-frequency content evident in the time courses can be attributed to the removal of high-power, low frequency content that is likely associated with cardiac activity, as evidenced by the differences in the power spectra before and after cleaning.

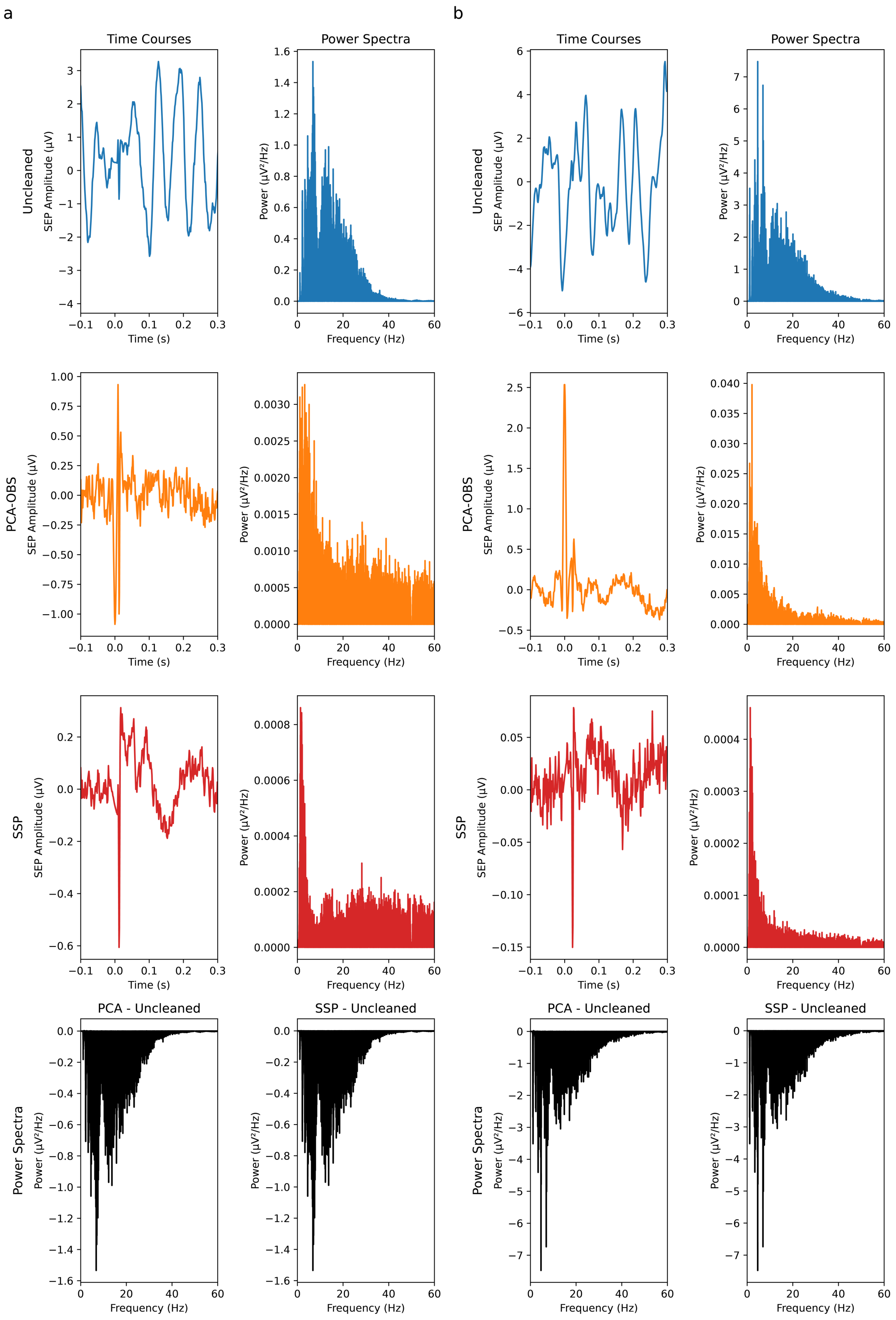


*Figure S5: Single participant (sub-020) depiction of the time courses and power spectra at low frequency of the data in the cervical spinal cord (a) and the lumbar spinal cord (b) without cleaning (top row), with PCA-OBS cleaning (second row) and SSP cleaning (third row). The bottom row depicts*

*the difference in the power spectra at low frequencies between the Uncleaned data and the two alternative cleaning methods. The much-reduced low frequency content in the time-courses demonstrating the somatosensory evoked potentials are explained by the much-reduced power of the low frequencies related to cardiac activity (mainly below 30Hz) as depicted in the power spectra. Please note that due to the difference in amplitude of low-frequency content in cleaned and Uncleaned data, the power spectra differences (black) are dominated by the Uncleaned data.*

### 5. Alternative ICA approaches

The inability of ICA to effectively remove the cardiac artefact while retaining spinal signals of interest was further investigated to determine whether alternative configurations could yield more satisfactory results. In the context of this study, the ICA pipeline was altered to determine if ICA could perform better with anteriorly re-referenced data (ICA-Anterior), or if ICA was applied separately to the cervical and lumbar spinal patches (ICA-Separated). As the results in Table S4 demonstrate, there are no consistent advantages to either ICA alternative tested.

*Table S4: Resulting residual intensity (RI), improved normalised power spectrum ratio (INPSR) of the cardiac artefact, and the signal-to-noise ratio (SNR) of the SEPs of the alternative configurations of ICA tested, computed across all participants.*

| | *RI* | *INPSR* | *SNR* |
|---|---|---|---|
| | | Cervical Spinal Cord | |
| *ICA* | 0.19% | 2.95E+90 | 3.30 |
| *ICA-Anterior* | 0.03% | 1.25E+95 | 2.89 |
| *ICA-Separated* | 0.12% | 8.50E+88 | 2.28 |
| | | Lumbar Spinal Cord | |
| *ICA* | 0.58% | 8.51E+93 | 2.09 |
| *ICA-Anterior* | 0.01% | 2.29E+92 | 2.48 |
| *ICA-Separated* | 0.07% | 4.96E+91 | 1.50 |

### 6. SEP time courses after the application of CCA

The plots depicted in Figure S6 demonstrate the similarities of the SEPs in both the cervical and lumbar spinal cord after the application of CCA. This means that even in the absence of pre-cleaning of the cardiac artefact, it is possible to obtain high quality SEPs for further analysis.

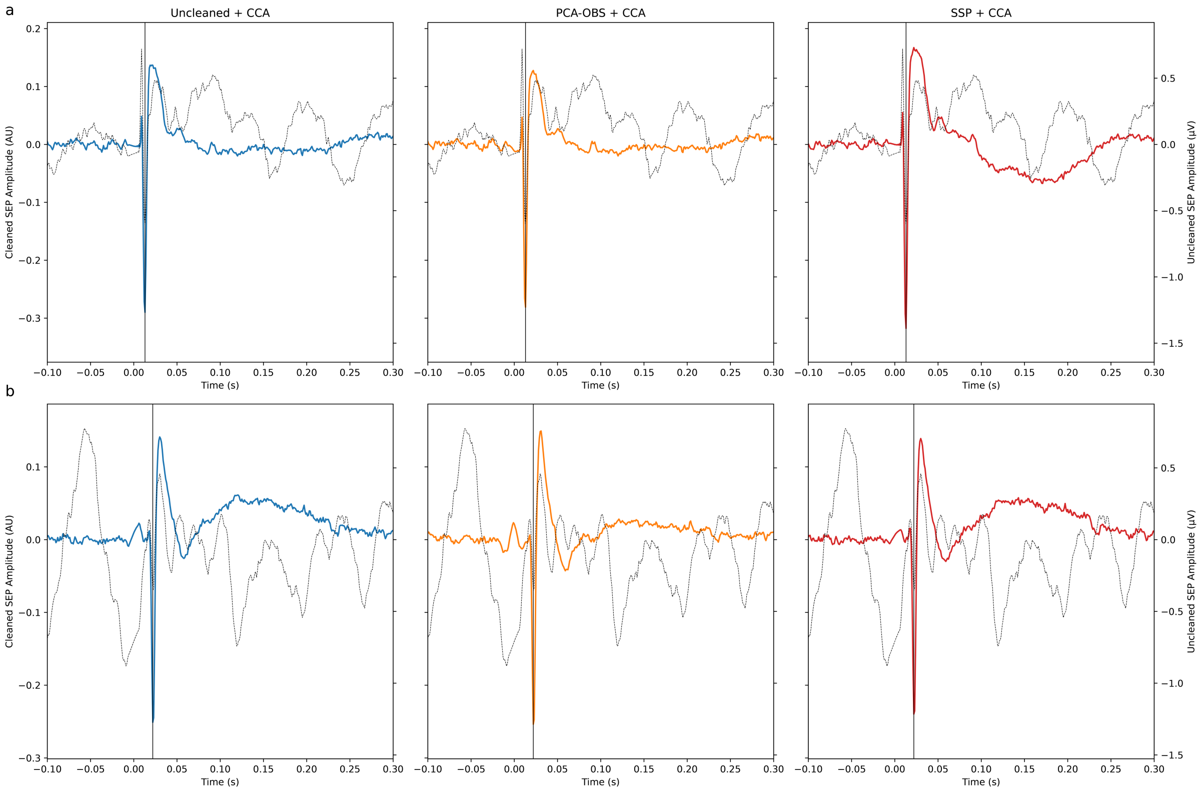


*Figure S6: Depiction of the grand average somatosensory evoked potential, for all participants and all trials, after additional processing via the application of CCA in the cervical (a) and lumbar (b) spinal cord. The left scale bar refers to the coloured traces in each plot after CCA has been applied. The right scale bar refers to the inset Uncleaned data (grey). The black line indicates the expected latency of the negative potential for median nerve stimulation (13ms) or tibial nerve stimulation (22ms).*

### 7. The ability of CCA to study single trial activity

A main advantage of CCA is its ability to enable the study of SEPs at the single trial level, as shown by Figure S7. Here, a single participant is depicted alongside negative peaks at the expected latency of SEPs in the cervical spinal cord in response to median nerve stimulation. Figure S7a shows the improvements that can be achieved in the robustness of single trial responses after cardiac artefact cleaning, with the best visual improvement seen in relation to data cleaned using SSP. Figure S7b demonstrates the impact of CCA in addition to cardiac artefact cleaning on the single-trial responses; here it is clear that robust single trial responses can be obtained using CCA, even in the absence of dedicated cleaning of the cardiac artefact.

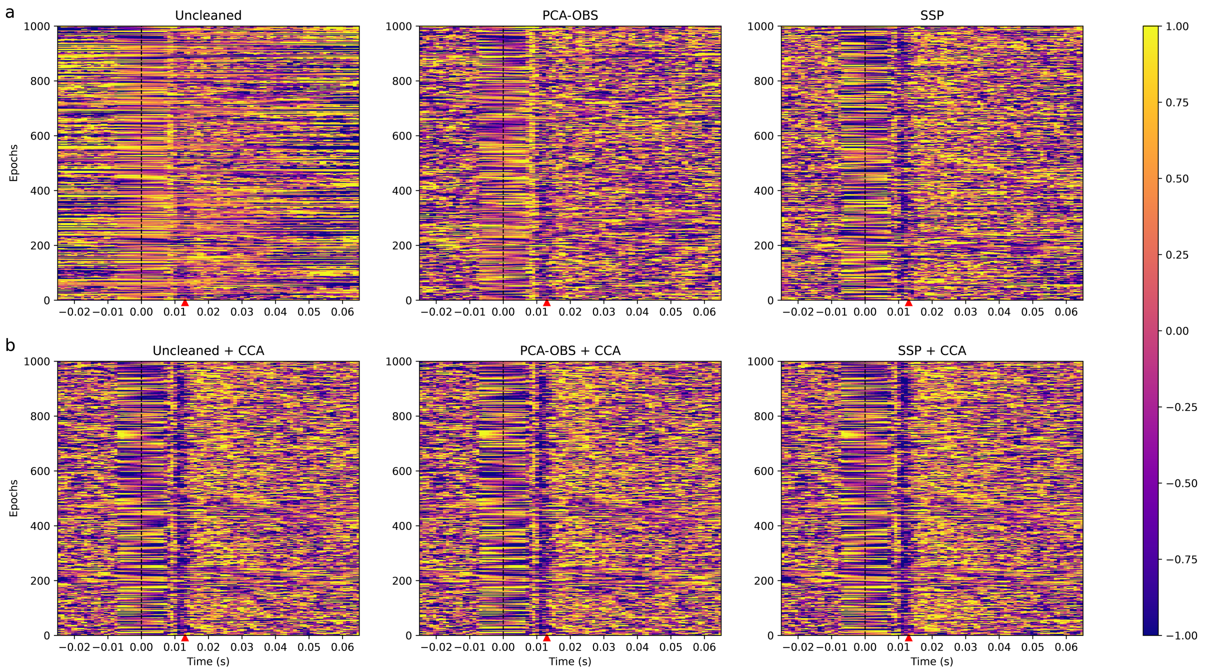


*Figure S7: Single trial plots for a single participant (sub-006) whereby each line in a plot is a different trial, created using 1/2 of the available trials in response to median nerve stimulation. (a) Shows the respective data prior to the application of CCA in cervical channel SC6, while (b) demonstrates the difference after CCA in the case of each cleaning regime for component 1. Each trial has undergone a z-score transformation to allow for comparison. The red triangle marks the expected latency of the SEP in each case.*
